# Fingerprints of high-dimensional coexistence in complex ecosystems

**DOI:** 10.1101/652230

**Authors:** Matthieu Barbier, Claire de Mazancourt, Michel Loreau, Guy Bunin

## Abstract

The coexistence of many competing species is a long-standing puzzle in ecology. Classic niche theory explains coexistence by trade-offs between a few essential species traits. Here we study an unexplored frontier of this theory: we assume that coexistence is intrinsically high-dimensional, arising from many traits and trade-offs at once. Species interactions then appear almost random, but their disorder hides a diffuse statistical structure: competitors that become successful start by subtly favoring each other, and partitioning their impacts on other species. We validate our quantitative predictions using data from grassland biodiversity experiments. We conclude that a high biodiversity can be attained through a pattern of collective organization that cannot be understood at the species level, but exhibits the fingerprint of high-dimensional interactions.

## Introduction

While species-rich ecosystems are often too complex for an exhaustive description, we can strive to understand them from two complementary directions. The first is to identify the role of particular species, traits and environmental factors, such as essential resources or keystone predators. The second approach is to search for collective phenomena that cannot be ascribed to any individual species, but emerge from their interplay. This latter notion has long appealed to theorists [1], but has so far received limited empirical support.

We claim that collective organization is indeed present in empirical ecological communities, and plays a demonstrable role in maintaining their biodiversity. Explaining how many species can coexist is a long-standing puzzle in ecology: various theories and experiments have brought forward the principle of competitive exclusion, whereby the best competitor should displace all others [5, 6]. Yet, strict dominance by one species appears, at most spatial and temporal scales, to be the exception rather than the rule in the natural world, and many communities host a remarkable diversity of competitors [7].

Over decades of ecological research, many partial solutions to this puzzle have been proposed, and most have been integrated into the overarching framework of niche theory [8, 9]. This framework suggests that we should identify *coexistence mechanisms*, particular trade-offs between species traits such as resource exploitation [10], defense against predators [11] and tolerance to environmental fluctuations [12, 13]. Through these trade-offs, strict order is imposed upon how species grow and interact (Fig. 1a), preventing any species from overwhelming its competitors.

**Figure 1:**
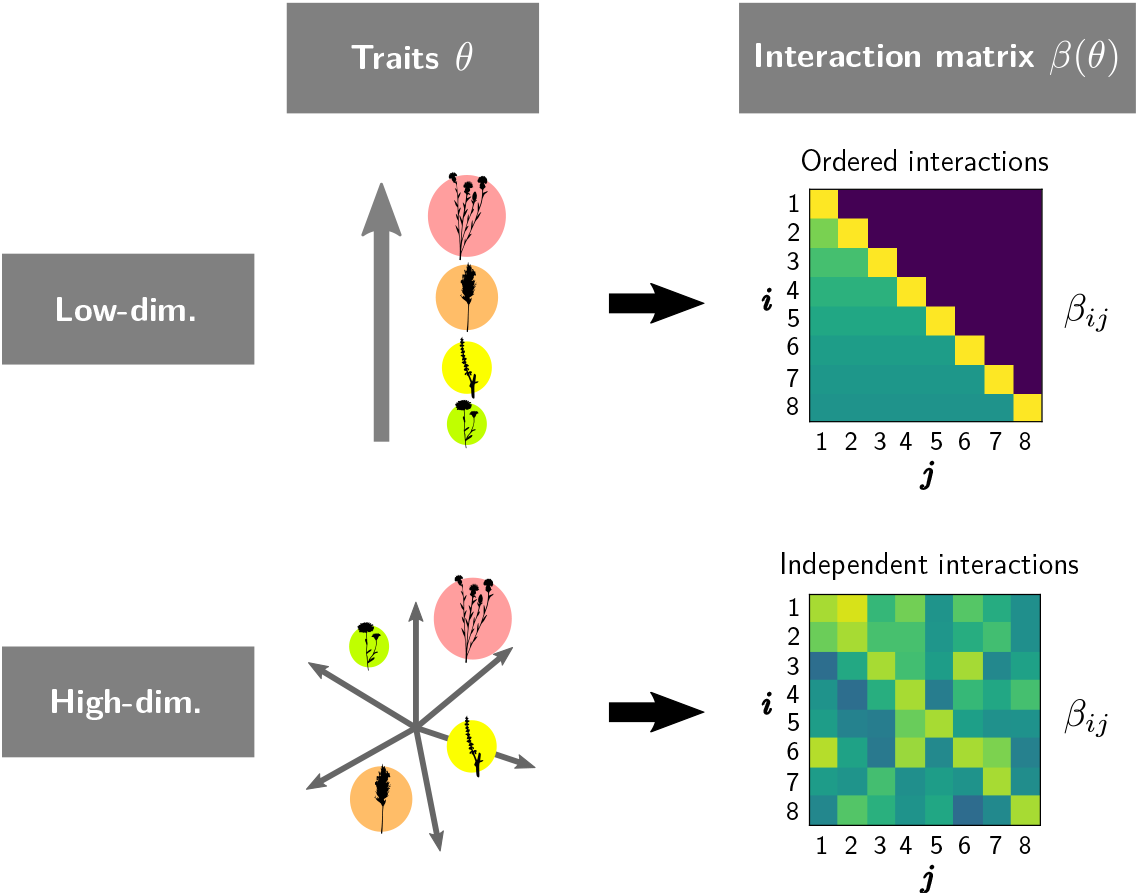
Dimensionality of interaction patterns. Assuming constant pairwise interactions, such as the Lotka-Volterra model in (2), the interaction network between *S* species can be represented by a *S* × *S* matrix (square box) where each element *β*_*ij*_ denotes the effect of species *j* on *η*_*i*_, the relative yield of species *i* defined in (1). **(a)** We assume that interactions *β* are entirely determined by underlying species traits and limiting factors *θ*, such as resources, pathogens or behaviors. In a low-dimensional trait space, each species is characterized by a few traits, therefore the interaction matrix can be ordered by trait values and displays a simple organization. **(b)** In a high-dimensional trait space, each interaction is a complex combination of factors, potentially unique and independent of other interactions. This can lead to a matrix without conspicuous order, often approximated by a random matrix [1, 2, 3, 4].

Niche theory has often been contrasted with a null model, the neutral theory of biodiversity [14], which predicts the behavior of communities of identical species with no coexistence mechanisms. In line with Clark [15], we propose a diametrically opposed null model: a community with many different coexistence mechanisms. Indeed, coexistence in highly diverse communities likely involves a large number of niches and trade-offs, some known and many unknown *a priori* [16]. The interaction between each pair of species may be determined by a unique combination of factors, precluding any simple and conspicuous (low-dimensional) order in the community [15]. As illustrated in Fig. 1, the interactions that emerge from a high-dimensional trait space appear almost independent of each other. Consequently, some ecological theories make the assumption that species interactions are essentially random [1] – a bold move, yet one that parallels major successes in physics and other fields [2, 3, 4]. Fully random interactions, however, do not allow many species to coexist [17, 18, 19]. The high biodiversity observed in many natural communities therefore implies some form of latent structure.

We predict that species coexistence can be achieved without any conspicuous order in the ecological network. Coexistence only requires a subtle statistical pattern, where “diffuse” correlations throughout the network bias how the most successful competitors interact with other species. We uncover this latent structure by an inferential approach shown in Fig. 2. We ask: if one samples many different interaction networks, and retains only those in which all species survive, what do the remaining networks have in common? Some may appear very structured, others almost random. Yet, we find that *most* of these networks exhibit the same statistical pattern, expressed in equations below. We derive this pattern from a simple probabilistic argument, explain in intuitive terms how it allows coexistence, and validate this pattern quantitatively in interaction networks inferred from biodiversity experiments.

**Figure 2:**
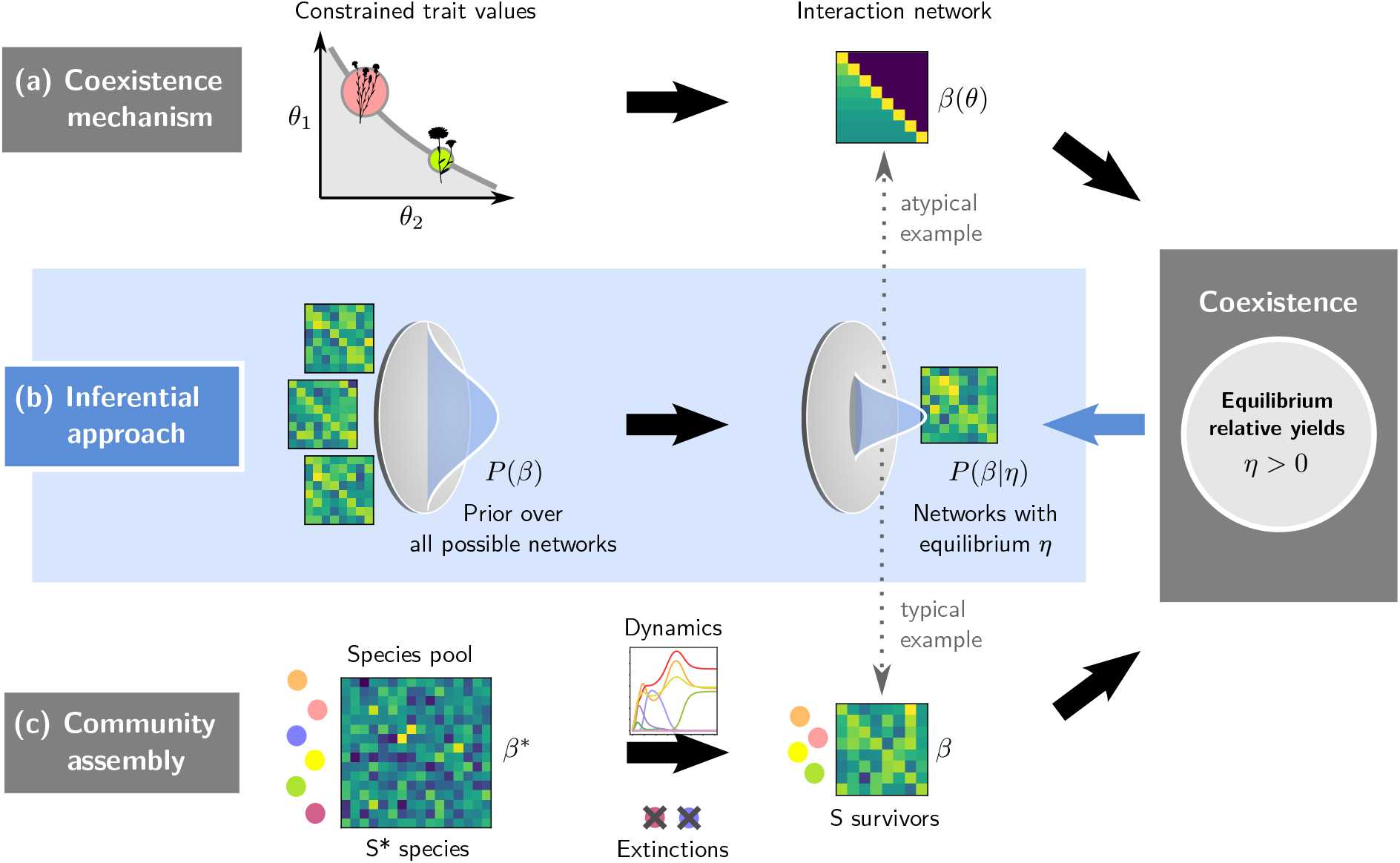
Three approaches to coexistence. All approaches are applicable to both low and high-dimensional settings (Fig. 1). **(a)** An equilibrium coexistence mechanism can be expressed as a specific pattern in the network of species interactions. To allow coexistence - i.e. an equilibrium where all species have positive relative yields *η* as defined in (1) – the traits *θ* which determine species interactions *β*(*θ*) must follow particular relations or constraints. We show here the classic low-dimensional example of a trade-off between two traits (competitive ability and colonization rate [20, 21], see Appendix B). **(b)** Our inferential approach reverses the direction of reasoning: given that we observe species coexisting at equilibrium in nature, we can infer the most likely distribution of interactions that could have caused this equilibrium, i.e. the posterior distribution *P* (*β|η*) given a prior *P*(*β*) over all possible interaction networks. If we choose a high-dimensional prior (independent interactions), communities drawn from the posterior distribution retain random-looking but correlated interactions, with simple and predictable statistical features: a “diffuse clique” pattern described below. **(c)** Community assembly involves a larger pool of *S*\* > *S* species, here with independent interactions. Through ecological dynamics, some of these species go extinct or cannot invade, leading to a smaller persistent group of *S* species whose interactions *β* have been dynamically selected to allow coexistence. **Dotted arrows:** Bunin [22] showed that the assembled community in (c) precisely follows the statistical properties of communities drawn from the posterior distribution in (b), if we assume independent interactions both for the inference prior and pool distribution. Low-dimensional structures, such as illustrated here in (a), can be seen as possible but very improbable matrices in this distribution, and can exhibit different patterns from our diffuse clique prediction. We can interpret the posterior distribution *P* (*β|η*) as a null model for coexistence in the presence of many biological mechanisms. The resulting diffuse clique pattern represents a form of collective organization, where coexistence arises, not from particular species traits, but from statistical biases distributed over all interactions.

## Results

### Theoretical pattern

We must first specify what we mean by interactions, and how they are estimated in real communities. Measuring species interactions is often difficult and prone to high uncertainty [23, 24, 25, 26], and most empirical settings only give us access to averaged properties. In particular, the total effect of interactions on one species *i* can be inferred from its *relative yield*

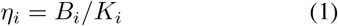

the ratio of its abundance *B*_*i*_ in a community to its maximum abundance *K*_*i*_ (known as carrying capacity) without competitors in the same environment [9]. We interpret species with higher *η* as *successful* competitors, as they benefit more in total (or suffer less) from their interactions with others. The simplest way to model these interactions is to assume a linear dependence between species’ relative yields

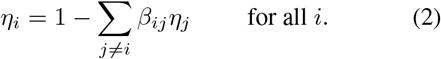

where *β*_*ij*_ is the competitive effect of species *j* on species *i* (which need not be symmetrical, i.e. it can differ from the effect of *i* on *j*). The relationship (2) has had some predictive success on data [27, 28]. It can also be derived from the classic Lotka-Volterra competition model, where it holds between coexisting species at equilibrium. The neutral theory of biodiversity [14] assumes *β*_*ij*_ = 1, while the existence of a pairwise co-existence mechanism for species *i* and *j* entails *β*_*ij*_ < 1.

Many different interaction networks (here, coefficients *β*_*ij*_) can generate the same equilibrium community. Observing the coexistence of *S* species conveys some information about their interactions, but not enough to fully determine them: the equations (2) impose *S* constraints, while there are *S*(*S* − 1) unknown interaction coefficients *β*_*ij*_. On the other hand, community-wide statistics, such as the expected strength of competition 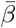, can reliably be deduced from the species’ relative yields [29] (see Appendix E).

We therefore adopt a probabilistic approach (Fig. 2), and ask what is the *most likely* community structure, i.e. the set of features most widely shared among the many possible solutions. We first define a prior distribution on the matrix elements, *P*(*β*), that can be adapted to our biological knowledge of a given community. For the experiments below, we simply assume that each coefficient *β*_*ij*_ is drawn independently from a normal distribution with mean 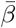. We then compute how this prior is modified once restricted to networks that admit the equilibrium *η*_*i*_ (Appendix B). Computing a posterior distribution given a prior and linear constraints (2) is a well-established problem in probability theory [30, 31].

We find that interactions *β*_*ij*_ should follow two statistical patterns that both admit intuitive interpretations (Fig. 3). First, competition must be biased to explain which species are successful or not. Here, success is relative to a baseline

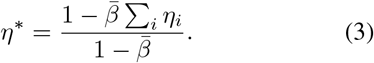

which is the relative yield that a species *would* achieve if all interaction strengths were equal to the prior mean (i.e. if hypothetically 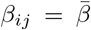). When *η*_*i*_ > *η*\*, we therefore expect that species *i* suffers less competition than the prior mean, and conversely if *η*_*i*_ < *η*\*. In our calculation, this appears in the conditional expectation of the competitive effect of *j* on *i* knowing their relative yields,

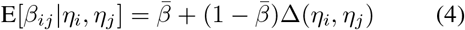

which deviates from the prior mean 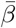 by a *competitive bias*

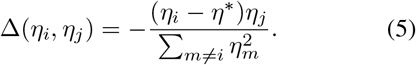

**Figure 3:**
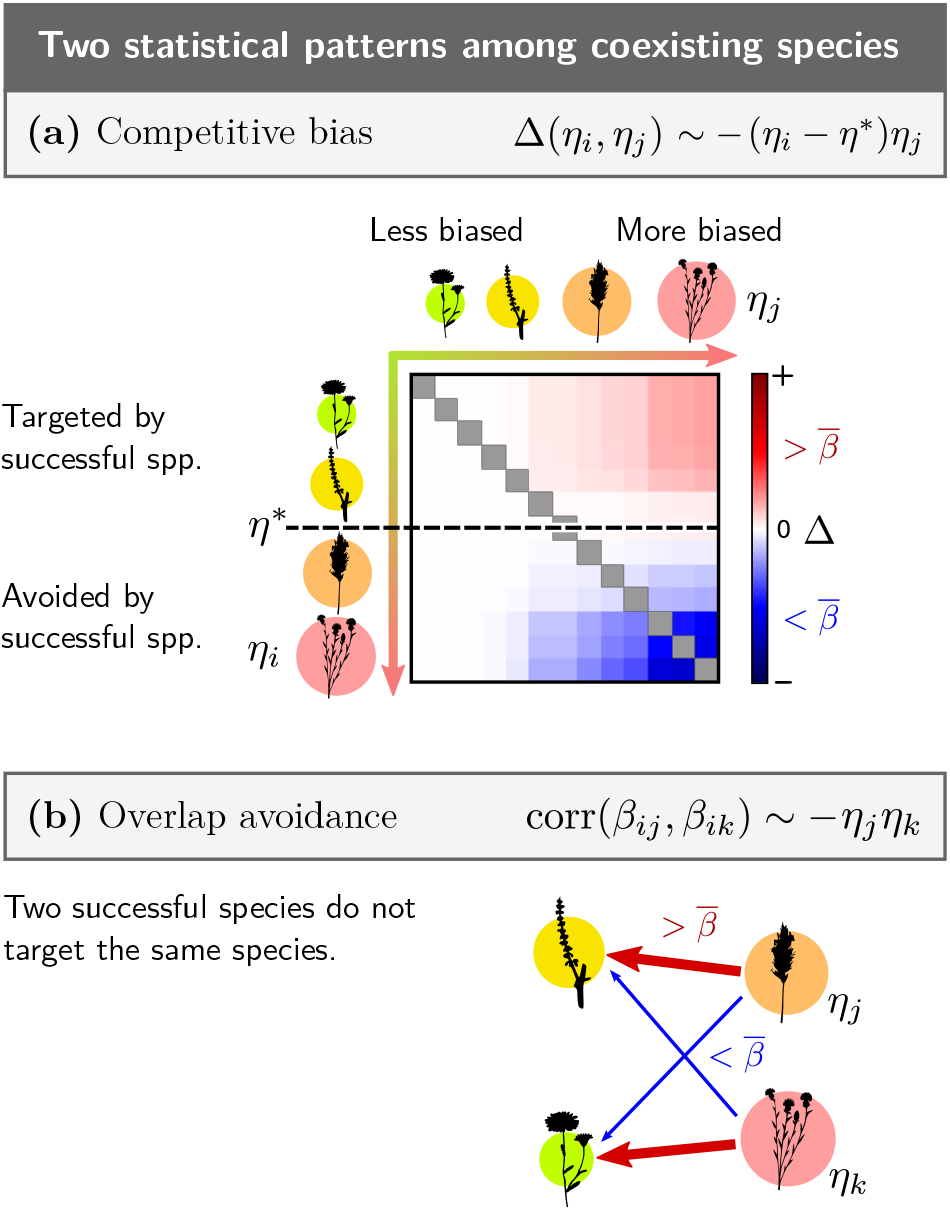
The diffuse clique structure is characterized by two statistical patterns in how successful species compete. Any typical interaction network drawn from the posterior distribution *P*(*β|η*) (see Fig. 2) will have similar statistical features. **(a)** Trend in the expectation of competition strength (4). If all species competed with equal strength 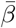, we would expect any given species *i* to achieve the relative yield *η*_*i*_ = *η*\* (3). Given how much *η*_*i*_ differs from this baseline, we can infer how interactions most likely deviate from 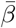. We show this deviation Δ(*η*_*i*_, *η*_*j*_) for simulated data, in matrix form: each element shows the bias in the interaction effect of species *j* (column) on species *i* (row). **Left to right:** an unsuccessful species (low *η*_*j*_) competes indiscriminately against others (white, Δ = 0), whereas a successful species (high *η*_*j*_) is a biased competitor. **Top to bottom:** species with *η*_*i*_ < *η*\* experience stronger competition from successful species (red, Δ > 0) whereas species with *η*_*i*_ > *η*\* experience weaker competition from them (blue, Δ < 0). Together, these biases indicate the existence of a “clique” of species that compete less against each other, and more against all others, thus achieving higher relative yield than the baseline *η*\*. **(b)** Correlation structure (6) between columns of the interaction matrix: the competitive effects of two successful species are anti-correlated, less likely to target the same species, whereas competition from unsuccessful species is indiscriminate.

We see that this bias is not evenly distributed among species. Competition coming from unsuccessful species (low *η*_*j*_) can be random without compromising the equilibrium. On the other hand, a species that is successful (high *η*_*j*_) is likely to compete less on average against other successful species, and to experience weaker competition from them (Fig. 3a).

The second pattern states that successful species *j* and *k* are less likely to compete against the same target *i* (Fig. 3b)

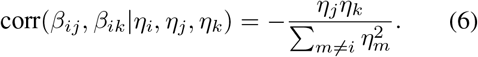

While the first pattern (4) determines the *expected* success of each species, the second pattern guarantees that each relative yield is *exactly* set to *η*_*i*_. We show in Appendix B that this correlation pattern prevents *η*_*i*_ from deviating from the value imposed by the competitive bias. Such deviations would likely drive some low-*η* species to extinction in a more random community.

### Empirical validation

We now present an empirical validation of these patterns on experimental data in Fig. 5 and Fig. 6. Grassland biodiversity experiments [33, 34, 35] provide an ideal testbed for inferring species interactions and mechanisms of coexistence. Each experiment contains a large number of plots in which plant species are assembled in varying numbers and combinations, out of a pool of *S* = 8 to 16 species depending on the experiment. Biomass in monoculture (single-species plots) provides an estimate of the species’ carrying capacities.

We first demonstrate the applicability of the equilibrium Lotka-Volterra model in Fig. 4. In the Wageningen grassland experiment especially, we observe a clear trend toward equilibrium (Appendix C). Pair-wise interactions almost all satisfy *β*_*ij*_ < 1, meaning that most species can coexist in pairs, with *E*[*β*] ≈ 0.4 and *std*(*β*) ≈ 0.2 showing a strong departure from the neutral hypothesis (*β*_*ij*_ = 1). Furthermore, these interactions *β*_*ij*_, inferred only from plots with 1 to 4 species, correctly predict relative yields in the full 8-species community (Fig 4a). Thus, the biodiversity in this experiment is well explained by our simple equilibrium model, without resorting to more complex explanations such as nonequilibrium coexistence [12] or higher-order interactions [36]. Following the method in [32], we show in Fig 4c and d that even in high-diversity experiments where all pairwise interactions cannot be inferred precisely, their statistics alone correctly predict the mean and variance of equilibirum relative yields.

**Figure 4:**
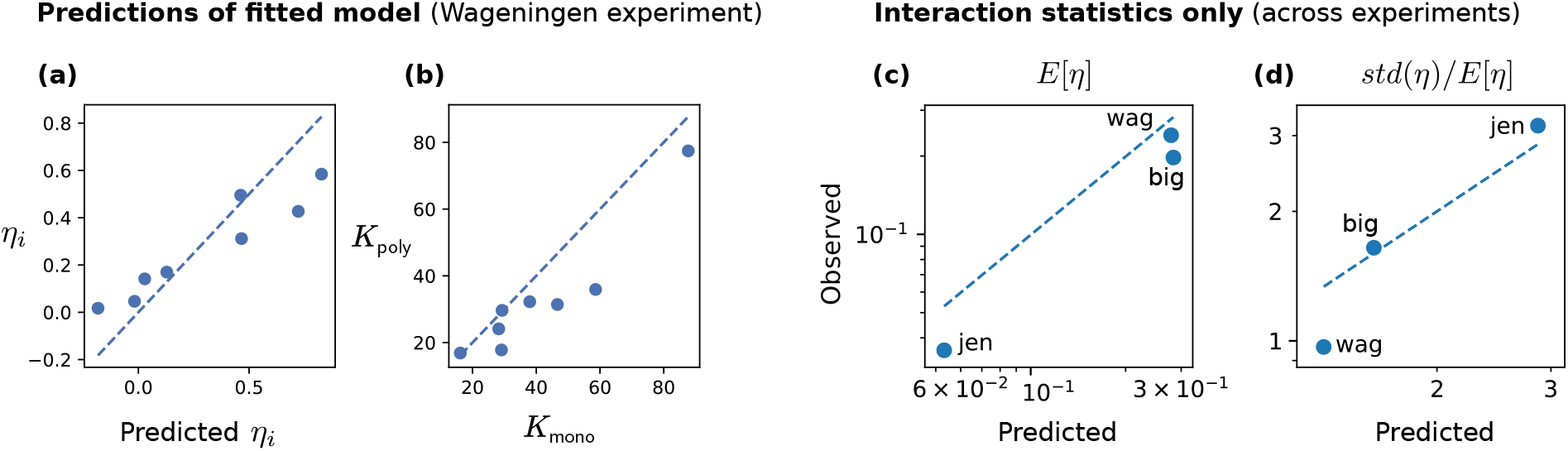
Validation of the Lotka-Volterra model. The consistency of the equilibrium description (2) was validated through a series of tests. For the Wageningen grassland experiment (with *S* = 8 species), we find accurate predictions for the full fitted Lotka-Volterra model (dashed lines show a 1:1 relationship between prediction and measurement). **(a)** We compare the equilibrium values of *η*_*i*_ = *B*_*i*_/*K*_*i*_ in the full-diversity plot (*S* = 8) to the prediction *η*_*i*_ = 1 − ∑_*j*_ *β*_*ij*_*η*_*j*_ where interactions *β*_*ij*_ were inferred from all partial compositions (plots with *S* < 8). **(b)** Carrying capacities *K*_*i*_ inferred as the intercept of the multilinear regression from polyculture plots (*S* > 1) agree with direct measurements in monocultures (*S* = 1). **(c)** and **(d)** Even when species are too numerous to allow a precise inference of all pairwise interactions, we can predict statistics of *η*_*i*_ using only the mean and variance of *β*_*ij*_ following the methodology in [32]. Across the Wageningen, Big Bio (16 species) and Jena (60 species) experiments, we find good agreement in predicted versus measured values for the mean in (c) and coefficient of variation in (d) of relative yields in the full-diversity plots.

**Figure 5:**
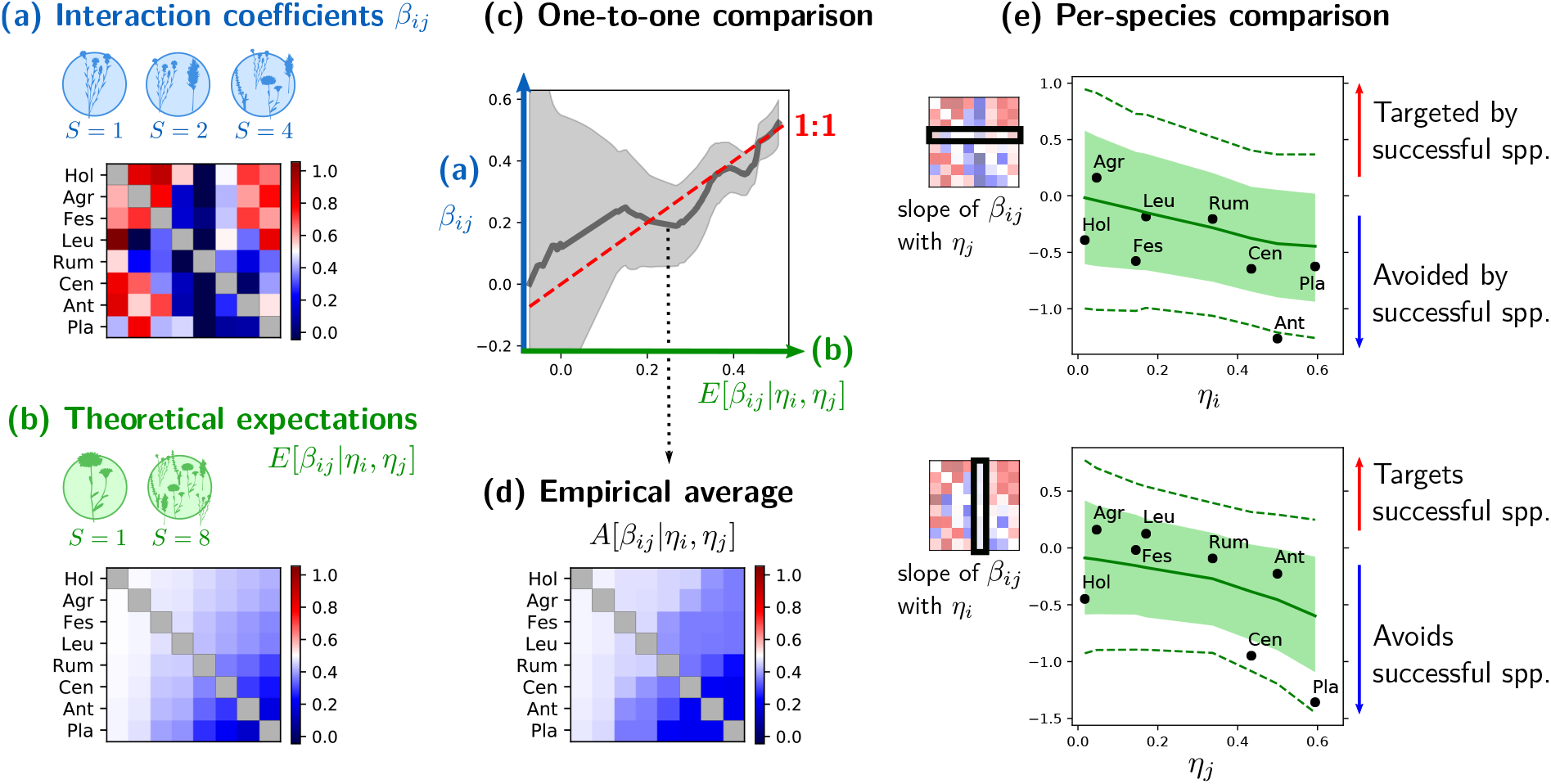
Diffuse structure in the Wageningen grassland experiment. **(a)** Using many species combinations (56 sets of *S* < 8 species), we can fit interaction coefficients *β*_*ij*_ in (2) by multilinear regression, and compute their mean *E*[*β*] ≈ 0.4. **(b)** Using the relative yields *η*_*i*_ in the full community (*S* = 8 species), we predict the theoretical pattern of expectations (4) for each interaction (Fig. 3a, here we see only negative biases in blue, indicating that all interactions are less competitive than the prior mean 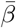). **(c-e)** In our diffuse pattern, individual coefficients are expected to exhibit a large spread around their expectation *E*[*β*_*ij*_|*η*_*i*_, *η*_*j*_], and must be binned to compute empirical statistics. **(c)** We group coefficients *β*_*ij*_ by their associated *η*_*i*_, *η*_*j*_ to compute the running average *A*[*β*_*ij*_|*η*_*i*_, *η*_*j*_] (grey curve, ±1 std. dev. in shaded area). Its proximity to the dashed 1:1 line is one of multiple metrics of theory-data agreement. **(d)** We show the empirical running average *A*[*β*_*ij*_|*η*_*i*_, *η*_*j*_] in matrix form (median value for each species pair *i, j*), to be compared with the theoretical prediction *E*[*β*_*ij*_|*η*_*i*_, *η*_*j*_] in (b). This is a particular way of smoothing the empirical matrix *β*_*ij*_ shown in (a), see Fig. S11 in SI. **(e)** We group coefficients by species, i.e. either by row or by column in the matrix *β*_*ij*_. We then compute how coefficients in row *i* (resp. column *j*) vary with *η*_*j*_ (resp. *η*_*j*_). Each solid point is the linear regression slope within a row or column. The empirical values and their spread around the trend are both in good agreement with our theoretical predictions, which also display some variance (±1SD within filled area, 90% CI between dashed lines) given that there are only *S* = 8 species.

**Figure 6:**
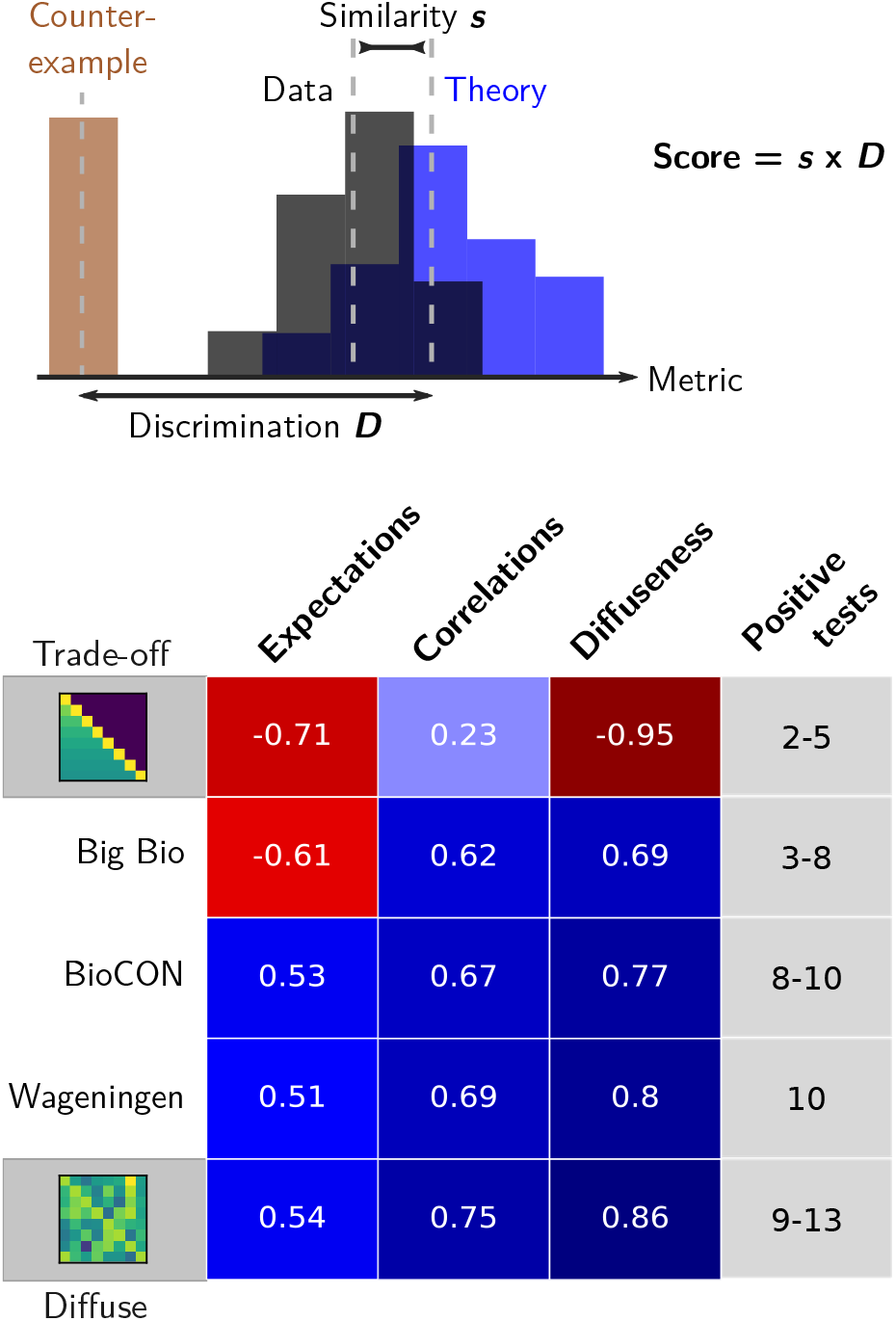
Cross-experiment validation of the diffuse clique structure. We compute metrics (defined in Appendix E) of agreement between data and theoretical predictions for conditional expectations *E*[*β*_*ij*_|*η*_*i*_, *η*_*j*_], correlations corr(*β*_*ij*_, *β*_*ik*_|*η*_*i*_, *η*_*j*_, *η*_*k*_), and diffuseness (whether interactions appear random). The first two metrics test the pattern shown in Fig. 3. We do not show each metric directly, because its predicted range is specific to each experiment. Instead, we assign a similarity score *s* = +1 if predicted and observed values differ by less than two standard deviations, −1 if they differ by more. We also compute a discrimination score *D* between 0 and 1 measuring how well the metric differentiates between our theory and a counter-example (see Materials and Methods). The product *s × D* is shown here. Other metrics were also tested, and the last column reports the number (or range across treatments) of positive results out of all 13 tests shown in Appendix F. The first and last row display simulated communities that constitute either counter-examples (low-dimensional trade-off patterns) or examples of our theory.

Our theory then predicts statistical trends in species interactions, given their relative yields in the full community plots^1^. The other plots, comprising subsets of the species pool, are used to fit the real interaction coefficients *β*_*ij*_ by a multilinear regression (2). From these fitted coefficients, we construct the empirical interaction matrix, and compare it to theoretically predicted expectations (4) and correlations (6).

We show in Fig. 5 the interaction matrix computed in the Wageningen grassland experiment [33], which supports our theory: individual coefficients *β*_*ij*_ display a random-looking spread, as we expect for a high-dimensional structure; yet we can group these coefficients in various ways to compute statistics, all of which concur with our predictions. We first bin interactions by their species’ relative yields. We can then compute the empirical running average *A*[*β*_*ij*_|*η*_*i*_, *η*_*j*_], which is close to a one-to-one relationship with the theoretical expectation *E*[*β*_*ij*_|*η*_*i*_, *η*_*j*_] (Fig. 5c). For more detailed insight, we group the coefficients by species: our theory (Fig. 3) predicts how the competition exerted and experienced by each species varies with the relative yields of competitors. Fig. 5e shows that empirical statistics agree with both predictions, with a spread around the trend comparable to what we expect theoretically for eight species.

This agreement between theory and data is precisely quantified for multiple experiments in Fig. 6. Our tests are designed to show that the empirical interactions appear too random to arise from a low-dimensional structure, yet exhibit the predicted pattern of means and correlations. We employ multiple metrics, each focusing on a different feature of our predictions, and we estimate how well each metric can discriminate between a positive example (a matrix generated according to our theory) and a negative one (here, a low-dimensional competition-colonization trade-off structure which does not follow our theory, see Fig. 2). Out of 13 tests detailed in Appendix F, we show three representative examples: two evaluate the predictions in Fig. 3 and one quantifies the diffuse (random-looking) character of the interactions. These tests taken together indicate moderate to good agreement for the Big Bio, BioCON and Wageningen experiments. The Wageningen experiment displays both the least noise in inferred coeffcients, and the most consistent and precise match to theory.

## Discussion

We have identified the most parsimonious way in which a complex network of ecological interactions can be organized so that all species coexist. We have shown that this pattern indeed occurs in plant interactions measured in several biodiversity experiments. This invites a new, more collective outlook on how to explain and predict species coexistence and biodiversity.

Our theory predicts that, when complex interactions arise from many different factors, ecological communities will most generally display a fuzzy “clique” of competitors that are both successful and less likely to compete strongly with each other, surrounded by unsuccessful species with random-looking interactions. This picture differs in multiple respects from classic explanations of coexistence. By imposing only the weakest possible constraints upon the many degrees of freedom in *β*_*ij*_, it allows individual interactions to take almost arbitrary values. It does not suppose a measurable segregation of species into distinct niches. It also represents a form of collective organization, where coexistence arises, not from particular species traits, but from statistical biases distributed over all interactions. Accordingly, this structure becomes increasingly likely to occur (although increasingly diffuse and subtle) in more diverse communities.

We stress that our results constitute a very stringent test for a theory of ecological coexistence. The fact that our simple multilinear model (2) correctly predicts coexistence (and even abundances) in various compositions is already a nontrivial result [26, 25, 29, 27, 28]. It suggests that more complex explanations, such as nonequilibrium coexistence [12] or higher-order interactions [36], are not required to explain the maintenance of biodiversity in these experiments. We go significantly further by demonstrating that the fitted interactions follow a theoretically-predicted pattern [22], fully determined by measurements without any adjustable parameter. We have shown that our test can tell apart this pattern from a low-dimensional trade-off mechanism [37]. To reduce inference biases, we use distinct abundance data to parameterize our theory and to infer the empirical interactions. We rule out these relationships being artefacts of our method, showing them to be violated in sparse and noisy data or counter-examples to our theory, see Fig. 6. Consequently, we believe that the experimental evidence is not merely suggestive, but strongly supportive of our claim.

In summary, we have introduced a novel way of thinking about species coexistence: not focusing on individual mechanisms, but predicting the universal statistical features that arise from combining many different coexistence mechanisms. We have quantitatively evidenced these features in real ecological communities. They reveal how *generic* an interaction network is: how statistically similar it is to most possible networks admitting the same equilibrium (Fig. 2). These features are not a necessary consequence of equilibrium coexistence: they can be violated in some communities, and these departures can hint at other biologically important low-dimensional mechanisms, such as simple trade-offs between ecological traits [37]. Instead, these features suggest the existence of a collective, intrinsically high-dimensional community organization. In such ecosystems, coexistence cannot be understood by analyzing particular species traits or environmental factors.

The classic alternative to niche theory has been the neutral theory of biodiversity [14], a null model that posits identical species without coexistence mechanisms. That model is empirically invalid here, as we observe heterogeneous and moderate species interactions that often allow pairwise coexistence. By contrast, our theory proposes a null model for complex communities with a large variety of coexistence mechanisms. By adapting our technique to different priors, it can be further applied to food webs, or other complex networks structured by ecological and evolutionary processes [38]. Our results imply the existence of a spectrum of possibilities between classic trait-based theories and complex network approaches, and provide a conclusive empirical test of collective organization in many-species coexistence.

## Materials and Methods

### Experimental data and interactions

We employ data from 3 grassland biodiversity experiments in Wageningen, Netherlands [33] and Cedar Creek, MN, USA (the Big Biodiversity [35] and BioCON [34] experiments). Each experiment uses a pool of species seeded or planted in various combinations, including some or all possible mono-cultures (*S* = 1 species), some partial compositions, and all species planted together. We removed the first two year for all experiments as they showed clear evidence of transient dynamics (Appendix C).

For each experiment, we first split monoculture (*S* = 1) data in two distinct sets, to be used separately, and computed the species’ carrying capacities *K*_*i*_ within each set. The first set of monocultures and all plots with 1 < *S* < *S*_*max*_ (where *S*_*max*_ is the experiment’s maximal plot diversity) were used to infer the interaction matrix *β*_*ij*_ using the hyperplane (multi-linear) least-squares fit proposed by Xiao et al [25] (see Appendix D). The second set of monocultures and the species abundance *B*_*i*_ in plots with *S* = *S*_*max*_ were then used compute the relative yields *η*_*i*_ = *B*_*i*_/*K*_*i*_ in the full community. All calculations were performed 250 times, using different boot-strapped sample means as values for *K*_*i*_ and *B*_*i*_. Each calculation led to a different set of *η*_*i*_, *β*_*ij*_ and 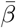 (see Appendix A for calculation details).

The multilinear fit of interactions [25] provided robust estimates of the effect of abundant species, but random fluctuations of *η*_*i*_ between plots could lead to very large (and highly variable) inferred per-capita effects for rare species with small *η*_*j*_. To avoid this issue, we first removed all species that had a reported abundance of 0 more than 90% of the time in plots where they had been planted, then removed coefficients *β*_*ij*_ whose variance between bootstrapped replicas was larger than the variance between coefficients in the matrix. This procedure retained all coefficients in the Wageningen experiment, but only around 60% of coefficients in the other experiments. Given that the Wageningen experiment gave more robust interaction values, we used it to assess our hypothesis that observed abundances are primarily determined by fixed pairwise species interactions according to (2). Fig. 4 shows strong empirical support for the hypothesis in this experiment. Interaction estimates from other experiments were less robust and might be affected by nonlinearity, transient dynamics, stochasticity and errors (Appendix D).

### Theoretical derivation

Any prior assumption about species interactions, such as the existence of coexistence mechanisms, can be expressed as a prior distribution *P*(*β*) over the space of all possible interaction networks *β* - i.e. a multivariate distribution over *S*(*S* − 1) coordinates which are the coefficients *β*_*ij*_ (excluding *β*_*ii*_ = 1). In Fig. 1, we translate the idea of an ecosystem with many different coexistence mechanisms into the prior that all coefficients are independent with mean 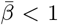 (allowing pairwise coexistence on average). In the following, we assume them to be normally distributed with variance var(*β*).

Even when pairs of species coexist, a fully random interaction network gives a vanishingly low probability of full coexistence in large communities [17, 18, 19]. If we see many species cohabit, we can infer the existence of non-random constraints on their competitive ties. These constraints can be expressed by a set of parameters *θ*, which determine the interaction coefficients *β*_*ij*_ (*θ*). We call any set of constraints a **coexistence mechanism** for our choice of dynamics if, given the matrix *β*(*θ*), these dynamics admit an equilibrium where all species survive, *η*_*i*_(*θ*) > 0 for all *i*.

In the inferential approach, we seemingly reverse the order of causality, as we infer the causes from the consequences (Fig. 2). The relative yields *η* are our clue to the underlying *θ*, which are conceived as hidden structural parameters of the interaction network. We show that we only need *S* parameters *θ*_*i*_ to guarantee, in Lotka-Volterra dynamics, that each species reaches a relative yield *η*_*i*_(*θ*) = *θ*_*i*_.

Thus, we define a posterior distribution *P*(*β|η*), but it is crucial to realize that the relative yields *η* are simply an **outcome**, and not a cause of this structure: successful (high-*η*) species do not exhibit biased competition by virtue of being successful; on the contrary, they are successful because they follow this interaction pattern. This arrow of causality is represented by our mathematical expression *η*_*i*_(*θ*) which is itself a result of dynamics on the network *β*(*θ*) parameterized by the values of *θ* (Fig. 3a).

To compute *P*(*β|η*), we impose the *S* equilibrium conditions (2), i.e. we restrict (project) the prior distribution *P*(*β*) to the hyperplane where ∑_*j*_ *β*_*ij*_*η*_*i*_ = 1 for all *i*. The projection of a multidimensional Gaussian distribution is still Gaussian, but with different statistics. Therefore, this posterior distribution *P* (*β|η*) is a new normal distribution over *S*(*S* − 1)-dimensional vectors *β*_*ij*_, entirely specified by its vector of means (4) and its covariance matrix (6), see Fig. 3. It represents the minimal deviation from our random prior that allows the observed community to coexist.

### Validation of theoretical predictions

We tested the two components of the diffuse clique pattern, as detailed in Appendix ES.

Starting with the pattern of means (4), we must compare the measured values of *β*_*ij*_ (hereafter *y*) to their theoretical expectation *E*[*β*_*ij*_|*η*_*i*_, *η*_*j*_] (hereafter *x*). To obtain an empirical estimate of the expectation for a single interaction coefficient, we performed a running average: for each point (*x, y*), we replaced its *y* coordinate by the average 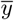 within a window centered on *x* and spanning 10% of the x-axis; we also measure the 90% CI over bootstrapped values (Fig. 3c). We then grouped all values 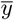 associated with the same species pair (*i, j*), took their median, and reconstructed an empirical matrix of expectations *B*_*ij*_ (shown in Fig. 3d). We also grouped coefficients by species *i* or *j* and performed regressions against *η*_*j*_ or *η*_*i*_ respectively (Fig. 3e), as well as the bi-linear regression of *β*_*ij*_ against *η*_*i*_*η*_*j*_ (the score computed from this metric is shown in Fig. 6 first column).

We proceeded similarly to test the pattern of correlations (6). Defining *d*_*ij*_ = *β*_*ij*_ − *E*[*β*_*ij*_|*η*_*i*_, *η*_*j*_] and the identity matrix 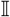, we computed for each species triplet (*i, j, k*) the value 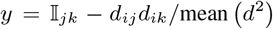, where the denominator is the sample mean over all pairs. We then did a regression of *y* against the prediction 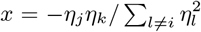 (Fig. 6 second column).

The diffuse nature of our pattern means that we expect the interaction matrix *β*_*ij*_ to appear almost random. True randomness is notoriously difficult to demonstrate [39], so we focused here on a more intuitive property: smoothness. The low-dimensional trade-off that we use as a counter-example creates a smoothly varying matrix *β*_*ij*_, whereas our diffuse pattern cannot. As a simple metric of smoothness, we measured differences between adjacent coefficients, and computed the fraction of these differences that are smaller than std(*β*) /2 (Fig. 6 third column).

For each of these metrics, we computed the distribution of values obtained from bootstrapped empirical matrices, and found its mean and variance 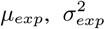. We also computed the distribution of values in matrices generated according to our theory 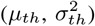, and in matrices generated with a competition-colonization trade-off 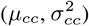, with the same equilibrium as the data (see Appendix B for details). In Fig. 6, we define similarity as

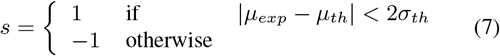

which states in a simple binary score whether observed values fall within the confidence interval (2 standard deviations) of predicted values. We define discrimination as

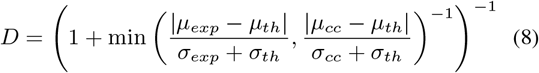

which quantifies how well a given metric can differentiate between patterns (*D* goes to 0 when standard deviations are significantly larger than the interval between means). The product *s × D* gives the scores shown in Fig. 6.

## Acknowledgments

We thank J.-F. Arnoldi, F. Isbell and Y. Zelnik for comments. Our gratitude goes to J. van Ruijven for sharing data from the Wageningen experiment.

## Data availability

This study brought together existing data that was obtained upon request (Wageningen biodiversity experiment data from J. van Ruijven [33]) and data that is publicly available (Big Bio http://www.cedarcreek.umn.edu/research/data, BioCON http://www.biocon.umn.edu/, Jena http://jenaexperiment.iee.uni-jena.de/.).

Data represented in Fig. 5 and 6 is available at https://github.com/mrcbarbier/diffuseclique.

## Footnotes

1 As shown in equations above, our theoretical predictions also depend on our prior for the mean interaction strength, 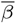. But this quantity is determined by the empirical average *E*[*β*] (see Appendix E). Hence, our theory has no freely-adjustable parameter.

