## Supplementary Appendices for "Fingerprints of high-dimensional coexistence in complex ecosystems"

### Supplementary Text

### A Overview

#### A.1 Model setting

Our approach for identifying the diffuse clique pattern can in principle be extended to arbitrarily complex interactions. But to present intuitive (and empirically successful) results, we choose here the simplest dynamical model of competition between $S$ species. For competitive interactions, it is classical to characterize the success of a species in competition by its *relative yield*

$$\eta_i = B_i/K_i \qquad \text{(Eq. S1)}$$

the ratio of its abundance $B_i$ in a community to its abundance without competitors $K_i$. We propose that dynamics are given by the generalized Lotka-Volterra equation[1]

$$\frac{d\eta_i}{dt} = r_i \eta_i \left(1 - \eta_i - \sum_{j \neq i}^{S} \beta_{ij} \eta_j \right) \qquad \text{(Eq. S2)}$$

where $r_i$ is the growth rate of species $i$, and $\beta_{ij}$ is the limiting effect of species $j$ on species $i$. This provides a minimal representation of interactions, expressed by one constant per species pair. At equilibrium, it imposes

$$\eta_i = 1 - \sum_{j \neq i} \beta_{ij} \eta_j \qquad \text{for all } i. \qquad \text{(Eq. S3)}$$

#### A.2 Summary of the approach

While we may not be able to infer precisely each interaction between coexisting species, the simple fact that these species coexist at equilibrium (with relative yield $\eta_i$) can allow us to infer the most likely statistical features of their interactions. **These features are not found exactly in any single community**, but most possible communities should be statistically close to this pattern. We clarify, following Fig. S1:

- Our theory defines expectations at the level of the *population* of possible interaction networks compatible with a given species' equilibrium $\eta$
- Each realized interaction network is a single sample from this population. Interactions observed in that network are not expected to follow exactly the theoretical expectations (in the same way that random variables drawn from a distribution are not expected to be exactly equal to the distribution mean).
- In addition to this true *variation* between networks (population variance), there is additional *error* in the inference process. Error should be minimized, and affects both the theoretical predictions (through error on their parameters $\eta$) and the measured interactions. By contrast, population variance applies only to measured interactions and cannot be minimized.

The approach detailed in this document can be summed up as the following:

1. **Setting** (Sec. A)
    - We observe a community of $S$ coexisting species, and measure each species' "competitive success" as its relative yield $\eta_i$ at equilibrium (its total biomass in the community divided by its total biomass in a similar environment without other species, which measures the cumulative effect of all interactions)

[1] See Sec.D.1 for a discussion, and in particular a justification of the use of relative yields $\eta_i$ rather than absolute biomasses.

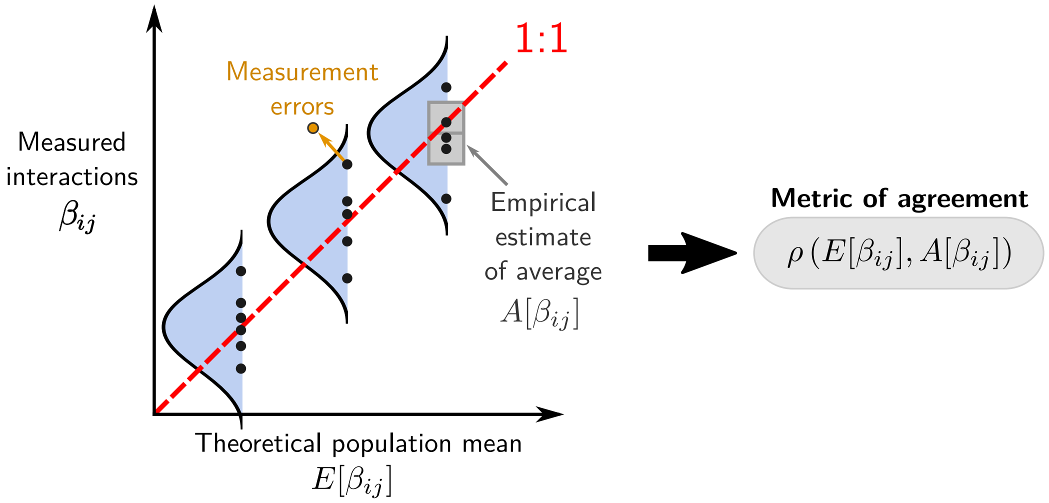

Figure S1: **Population statistics and sample measurements.** We predict that each interaction coefficient $\beta_{ij}$ is drawn from a distribution with an expectation $E[\beta_{ij}|\eta_i, \eta_j]$, biased by the relative yield of species $i$ and $j$. However, we usually do not expect a large correlation coefficient between individual coefficients $\beta_{ij}$ and predicted expectation $E[\beta_{ij}]$: even in synthetic data generated from our theory, the distribution's variance along the y-axis can be very large (potentially much larger than the range spanned by the expectation along the x-axis). Instead, we test the pattern by computing an empirical average $A[\beta_{ij}|\eta_i, \eta_j]$ (binning together multiple coefficients $\beta_{ij}$ that have almost the same expectation). This empirical average $A[\beta_{ij}]$ is the estimate that should be compared to the theoretical means $E[\beta_{ij}]$, using the correlative metric $\rho$ defined in (Eq. S36). We also note that while measurement errors can displace points in both dimensions, population variance only concerns the y-axis; hence the two axes cannot be treated equally (the averaging over population variance, going from $\beta_{ij}$ to $A[\beta_{ij}]$, should be distinguished from the treatment of errors in $\beta_{ij}$ itself or in the prediction of $E[\beta_{ij}]$ from $\eta_i$,$\eta_j$). The reasoning developped here for $E[\beta_{ij}]$ also applies to the other component of our statistical pattern: the predicted correlation structure $\mathrm{corr}(\beta_{ij}, \beta_{ik}|\eta_i, \eta_j, \eta_k)$ and corresponding measurements on pairs of interactions.

- Many possible interaction networks $\beta_{ij}$ (i.e. different community structures) could lead to the same relative yields $\eta_i$ for these coexisting species. There is too little information to specify each interaction coefficient $\beta_{ij}$, since there are $S(S-1)$ such coefficients, and only $S$ constraints (the values that $\eta_i$ should take when all species coexist together).
- But out of all the possible networks that are compatible with the same observed $\eta_i$, most of these networks will exhibit a statistical pattern in their coefficients. We derive theoretically this pattern and show that it has two components: a bias in the mean of interactions $\beta_{ij}$, and another in the correlations of pairs of interactions $\beta_{ij}$ and $\beta_{ik}$.

2. **Defining the pattern on means** (Sec. B)

   - We first characterize the *population mean* or *expectation* $E[\beta_{ij}|\eta_i, \eta_j]$. For a given pair of species $(i, j)$, each possible network will have a different value of $\beta_{ij}$, and this value can vary broadly between networks. But on average, this vast range of possibilities for a single coefficient $\beta_{ij}$ will be *biased*: the distribution of possible interaction coefficients for species pair $(i, j)$ should depend on their observed relative yields $(\eta_i, \eta_j)$.
   - As we will show, this bias follows a very intuitive pattern. On the receiving end $i$ of the interaction, less successful (low $\eta_i$) species should experience more negative interactions, while successful species should experience less negative (or more positive) interactions. On the giving end $j$, this bias is not uniform: only successful species (high $\eta_j$) need to be biased in who they affect. Finally, the slope of the bias is determined collectively, in a way that involves all species in the community.

3. **Testing the pattern on means** (Sec. D and E)

   - To test this pattern, we need an independent way of estimating the true interaction network $\beta_{ij}$ in a community. In competition experiments, we can measure the biomass $B_i^{(w)}$ of species $i$ in many treatments (e.g. plots of land with different combinations of the same plant species) $w$ with $S(w) = 1, 2, 4, ...$ species in them. Then, we may have enough information to fit each interaction coefficient $\beta_{ij}$ in that community (assuming that interactions are fixed pairwise coefficients that do not change between treatments).
   - We use all treatments with $S(w) < S_{\max}$ to get an empirical estimate of $\beta_{ij}$, and the treatment $S = S_{\max}$ to have the $\eta_i$ that we will use to compute predictions for the clique pattern $E[\beta_{ij}|\eta_i, \eta_j]$.
   - **We do not expect a strong correlation between individual empirical $\beta_{ij}$ and predicted $E[\beta_{ij}|\eta_i, \eta_j]$**. Indeed, as noted above, a given coefficient $\beta_{ij}$ in a given interaction network can have very large variation around the expectation that is predicted over all potential networks. This is not an error in the inference process: **this variance is part of our predictions**, and it is the reason why we call it a *diffuse* clique pattern – if we did not know precisely where to look, we might not be able to discern any pattern.
   - **To test the pattern, we must construct empirical averages to compare to the theoretical expectations $E[\beta_{ij}|\eta_i, \eta_j]$.** There are multiple ways of binning together different coefficients $\beta_{ij}$ to compute their statistics
     - We can bin coefficients $\beta_{ij}$ by species $i$ or species $j$, or by $\eta_i$ or $\eta_j$, and the compute the mean interaction within each bin, and the linear regression slope within the bin (the slope with $\eta_i$ when binned by species $j$, and with $\eta_j$ when binned by species $i$
     - We can bin together different coefficients $\beta_{ij}$, associated with different species pairs $(i, j)$, but with the same theoretical expectation. We then take the average $A[\beta_{ij}]$ over the bin, and our measure of success is the distance between $A[\beta_{ij}]$ and the theoretical expectation $E[\beta_{ij}]$ for the same bin.

4. **Defining and testing the pattern on correlations** (Sec. D and E)

- The same reasoning applies to correlations in the interactions, $\mathrm{corr}(\beta_{ij}, \beta_{ik}|\eta_i, \eta_j, \eta_k)$. We predict theoretically that there should be correlations between all the possible values that interactions $\beta_{ij}$ and $\beta_{ik}$ take in many different networks compatible with the same relative yields $\eta$.
- These correlations also have an intuitive interpretation: $\beta_{ij}$ and $\beta_{ik}$ are anti-correlated for successful species $j$ and $k$, indicating that two successful species will tend not to compete strongly against the same victim $i$. But once again, this only holds statistically and with large variation between possible communities. This is, therefore, a diffuse pattern of avoidance or partitioning in who competes against whom.
- To test this pattern in the empirically measured network $\beta_{ij}$, we first measure $d_{ij} = \beta_{ij} - E[\beta_{ij}|\eta_i, \eta_j]$, using the theoretical expectation defined above, and use it to construct $y_{ijk} = d_{ij} d_{ik}/\mathrm{var}(d)$ for each triplet of species $(i, j, k)$. Here $\mathrm{var}(d)$ is the variance of the entire set of values of $\Delta$, which is predicted to be independent of the identity of the species.
- Once again, **we do not expect a strong relationship** between the empirical $y_{ijk}$ in one network and the theoretical $\mathrm{corr}(\beta_{ij}, \beta_{ik}|\eta_i, \eta_j, \eta_k)$ across all possible networks. Each network is expected to vary very significantly around the trend.
- To identify this trend, **we must again perform empirical binned averages** (as for the pattern on means, see above)
  - we group $y_{ijk}$ by species $i$, and compute their mean and their slope with $\eta_j \eta_k$
  - we group $(i, j, k)$ triplets with the same theoretical expectation $\mathrm{corr}(\beta_{ij}, \beta_{ik}|\eta_i, \eta_j, \eta_k)$, and average their coefficients to get the empirical average $A[y_{ijk}|\eta_i, \eta_j, \eta_k]$.

5. Using these metrics (for the pattern of means, and for the pattern of correlations), we quantify how much our proposed diffuse community structure can be detected in real networks from coexistence experiments. We compare these measurements to those obtained in simulated networks, generated either according to our theory, or according to one possible competing hypothesis (a competition-colonization trade-off mechanism).

#### A.3 Summary of the analysis pipeline

For convenience, we provide here a high-level technical summary of the analysis pipeline employed to obtain the results presented in the main text, skipping the details and justifications found in later sections. We simply note that, because of the distinction shown in Fig. S1 between **population variance** (on the y-axis) and **measurement error** (on both the x- and y- axes), we adopt a bootstrap-based approach to errors: in short, we generate many bootstrap estimates of the true value of each inferred variable, then treat all these estimates as if they now had zero error. When we confront pairs of variables as in Fig. S1, we can therefore bin these points by their x-coordinate when estimating the average y-coordinate (i.e. averaging out the population variance) as if these coordinates were exact.

Our pipeline for each experiment:

1. Format data in tuples (Plot, Composition, Date, Species, Biomass), where Composition is the list of all the species planted in the plot.

2. Drop the first two years, which are noticeably out of equilibrium, see Sec. C.3.

3. Create a table **Biomasses** where, for each pair (Composition, Species), all Biomass data are regrouped into a vector $\vec{B}(c, s)$, dropping all information about Date and Plot.

4. From this table **Biomasses**, generate many tables of bootstrapped estimates $\mathbf{BiomassAvg}_n$ ($n = 1 \dots N_{replica}$) where

- each vector of biomasses $\vec{B}(c,s)$ of length $L_B$ is replaced by a vector $\vec{b}(c,s)$ of the same length $L_B$
- each element of the vector $\vec{b}(c,s)$ is computed as the average of $L_B$ numbers drawn from $\vec{B}(c,s)$ with replacement (bootstrapped average)
- hence, the standard deviation in each vector $\vec{b}(c,s)$ now approximates the standard error on the mean of the corresponding $\vec{B}(c,s)$ (in other words, variance between elements of $\vec{b}$ is $1/L_B$ of the variance between elements of $\vec{B}$)

5. Split each table $\mathbf{BiomassAvg}_n$ into two subsets:
   - the set **A** containing all compositions with $S < S_{\max}$ species, where $S_{\max}$ is the maximal number of species in a composition, and half of the points (selected at random) of each vector $\vec{B}$ associated with a *monoculture* (i.e. a composition including a single species)
   - the set **B**, containing all compositions with $S = S_{\max}$ species and the other half of the points of each monoculture vector.
6. For each table $\mathbf{BiomassAvg}_n$:
   - use the set **A** to infer an empirical interaction matrix $\beta_{ij}$ by the hyperplane regression described in Sec. D.2
   - use the set **B** to compute relative yields for each species $i$, $\eta_i = B_i/K_i$ where $B_i$ and $K_i$ are the average over the vectors $\vec{b}$ associated with the abundance of species $i$ in the full community and in monoculture,
   - use the $\eta_i$ from set **B** to parameterize the theoretical predictions for $\bar{\beta}$ (see Sec. E.3), $E[\beta_{ij}|\eta_i,\eta_j]$ and $\mathrm{corr}(\beta_{ij},\beta_{ik}|\eta_i,\eta_j,\eta_k)$.
7. Randomly draw integers $n$ and $n'$ and pair the set **A** from table $\mathbf{BiomassAvg}_n$ and the set **B** from table $\mathbf{BiomassAvg}_{n'}$, until all sets are paired.
   - Pair together each coefficient $\beta_{ij}$ from the set **A** of table $n$ with the corresponding prediction $E[\beta_{ij}]$ obtained from the set B of table $n'$
   - Use the coefficients $\beta_{ij}$ from the set **A** of table $n$ and the $E[\beta_{ij}]$ from the set **B** of table $n'$ together for two pairs of species $(i,j)$ and $(i,k)$ to compute $y_{ijk}$ as defined in the previous section
   - Pair together each $y_{ijk}$ with the corresponding prediction $\mathrm{corr}(\beta_{ij},\beta_{ik}|\eta_i,\eta_j,\eta_k)$ obtained from the set **B** of table $n'$.
8. Pool together all tables $\mathbf{BiomassAvg}_n$, and for each of the two sets of observables (i.e. $z$ can denote either $\beta_{ij}$ or $y_{ijk}$):
   - take the points $(x,y) = (E[z],z)$ (where $E[z]$ is the corresponding prediction) from all species and all tables
   - perform a running average, i.e. replace each point $(x,y)$ by $(x,y') = (E[z],A[z])$ where $A[z]$ is the average of all the values of $z$ included within a window centered on $x = E[z]$ and whose width is given by 10% of the full $x$-axis.
   - bin $x$ and $y$ by species, and compute within each bin the average coefficient and its linear regression slope against the $\eta$ of other species, to obtian
   - estimate the standard error on $A[z]$ as $\mathrm{std}(z)\,/\sqrt{\#(z)/N_{replica}}$ with the standard deviation of $z$ within the window, $\#(z)$ is the number of points within the window, and $\#(z)/N_{replica}$ is thus the average number of points within the window belonging to a *single* bootstrapped replica (this estimation thus becomes asymptotically independent from $N_{replica}$)

- Compute the agreement between the predicted expectations $E[z]$ and measured averages $A[z]$ for all $z$ in the dataset as

$$\rho(z) = \frac{\text{cov}(E[z], A[z])}{\max\left(\text{var}(E[z]), \text{var}(A[z])\right)} \qquad \text{(Eq. S4)}$$

  This definition ensures that $\rho$ coincides with the Pearson correlation coefficient when $A[z]$ and $E[z]$ have the same variance, but is reduced when their variances differ, so as to capture quantitative agreement in the slope and not only correlation.

- For each metric, compare this score to the distribution of scores obtained by applying the same procedure to simulated matrices (instead of the empirical matrix) with the same equilibrium $\eta_i$:
  - 500 matrices generated according to our pattern (Sec. B.4) with the same statistics $\langle\beta\rangle$ and $\text{var}(\beta)$ as the empirical matrix
  - 500 matrices generated according to the competition-colonization tradeoff pattern ((Sec. B.6)

  We can thus estimate whether the score of the empirical matrix falls within the range expected for our pattern given the abundances $\eta_i$ and the statistics $\langle\beta\rangle$ and $\text{var}(\beta)$, and whether this score allows us to discriminate a counter-example such as a simple tradeoff pattern.

9. For visualization purposes, construct matrices from $A[\beta_{ij}]$ and $\text{median}(E[\beta_{ij}])$ (the median is taken over results obtained for different tables $\mathbf{BiomassAvg}_n$) for each species pair $(i, j)$, and compare them to see the qualitative trend in each.

### B Diffuse clique pattern

#### B.1 Context: Coexistence mechanisms and hidden structural parameters in interaction networks

A fully random interaction network gives a vanishingly low probability of coexistence in large communities [**?**, **?**, **?**]. If we see many species cohabit, we can infer the existence of non-random constraints on their competitive ties. These constraints can be expressed by a set of parameters $\boldsymbol{\theta}$, from which we compute the interaction coefficients $\beta_{ij}(\boldsymbol{\theta})$. We call these constraints a **coexistence mechanism** for the dynamics (Eq. S2) if, given the matrix $\boldsymbol{\beta}(\boldsymbol{\theta})$, these dynamics admit an equilibrium where all species survive, $\eta_i(\boldsymbol{\theta}) > 0$ for all $i$.

In the inferential approach, we seemingly reverse the order of causality, as we infer the causes from the consequences. The relative yields $\boldsymbol{\eta}$ are our clue to the underlying $\boldsymbol{\theta}$, which are conceived as hidden structural parameters of the interaction network. We show that we only need $S$ parameters $\theta_i$ to guarantee, in Lotka-Volterra dynamics, that each species reaches a relative yield $\eta_i(\boldsymbol{\theta}) = \theta_i$.

Thus, we write $P(\beta|\eta)$, but it is crucial to realize that the relative yields $\boldsymbol{\eta}$ are simply an **outcome**, and not a cause of this structure: successful (high-$\eta$) species do not exhibit biased competition by virtue of being successful; on the contrary, they are successful because they follow this interaction pattern. This arrow of causality is represented by our mathematical expression $\eta_i(\boldsymbol{\theta})$ which is itself a result of dynamics on the network $\boldsymbol{\beta}(\boldsymbol{\theta})$ parameterized by the values of $\boldsymbol{\theta}$ (Fig. S2a).

#### B.2 Description

The diffuse clique pattern is the minimal set of constraints that allows coexistence, making no further assumption about species traits and interactions. Rather than a deterministic relationship $\boldsymbol{\beta}(\boldsymbol{\theta})$, we search for a distribution $P(\boldsymbol{\beta}|\boldsymbol{\theta})$ that is as close as possible to our prior $P(\boldsymbol{\beta})$ while still ensuring coexistence.

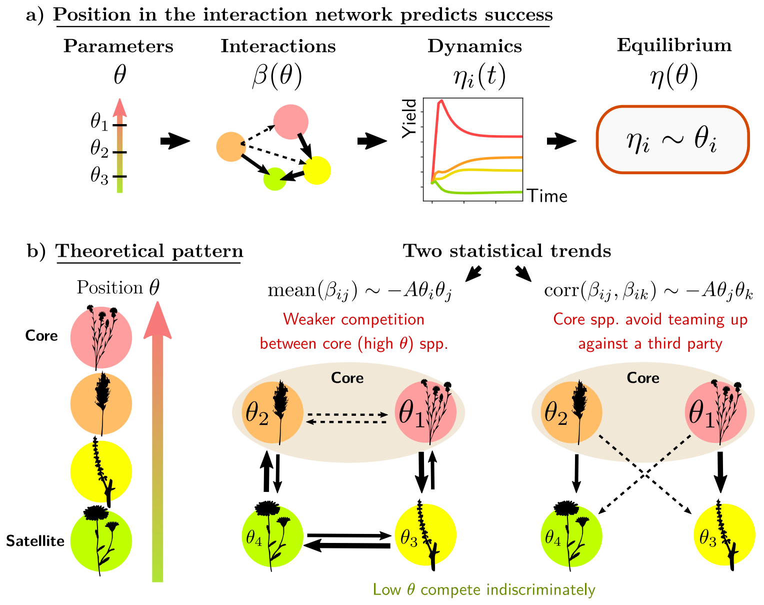

Figure S2: Structure of the diffuse clique pattern expressed in terms of the hidden structural parameters $\boldsymbol{\theta}$ proposed in Sec. B.1. **a)** The relative yields $\boldsymbol{\eta}$ are a consequence of the dynamics in a community whose interactions are determined by a set of parameters $\boldsymbol{\theta}$. Therefore, $\boldsymbol{\eta}$ can be used as a clue to infer these interaction parameters. This inference is especially simple in the Lotka-Volterra model, where the prediction is that $\eta_i(\boldsymbol{\theta}) = \theta_i$ at equilibrium, independently of other $\theta_j$. In other words, the "network position" $\theta_i$ that parameterizes the distribution of interactions of species $i$ will also determine exacty and uniquely its relative yield at equilibrium $\eta_i$. **b)** Core species with high $\theta$ tend to avoid competing with each other, which allows them to be more successful on average. In turn, they allow other species to survive, by distributing their competitive effect selectively, so as to avoid teaming up against the same species.

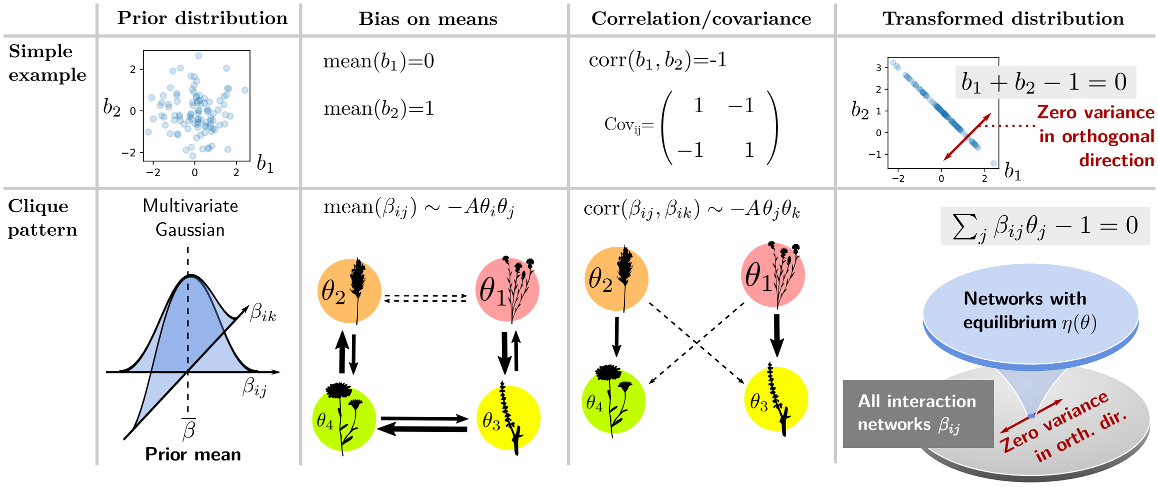

Figure S3: Illustration of how the biases in the means and correlations of interactions $\beta_{ij}$ suffice to ensure precisely the satisfaction of constraints (Eq. S5), and therefore, how they determine the relative yields $\boldsymbol{\eta}(\boldsymbol{\theta})$ at equilibrium. We first show the same ideas in a simpler example, with only two random variables, demonstrating how we can ensure that these variables $b_1$ and $b_2$ can both be drawn at random such that a constraint $b_1 + b_2 - 1 = 0$ is always satisfied. In that case, the points $(b_1, b_2)$ all fall on a line (since there are two variables and one constraint, there is only one degree of freedom left). The bias in the means determines the location of this line in the plane, and the correlation structure determines the direction of the line: it guarantees that the constraint is satisfied *exactly* by cancelling all variation in the orthogonal direction. In simpler terms, imposing a perfect anticorrelation of $b_1$ and $b_2$ guarantees that their sum never changes value. Likewise, imposing a correlation pattern upon the $\beta_{ij}$ can ensure that their weighted sum $\sum_j \theta_j \beta_{ij}$ is always set to 1.

To provide an intuitive understanding of this pattern, we use here the simple Lotka-Volterra model to derive explicit formulas, but we will show in Sec. E that its qualitative features are robust in empirical data. Given a prior on the interaction distribution $P(\boldsymbol{\beta})$, we can compute the conditional distribution $P(\boldsymbol{\beta}|\boldsymbol{\theta})$ that satisfies the following linear constraints:

$$\sum_j \beta_{ij}\theta_j = 1 \qquad \text{for all } i, \qquad \beta_{ii} = 1. \tag{Eq. S5}$$

When these $S$ equations hold, it is easy to check in (Eq. S2) that Lotka-Voltera dynamics admit an equilibrium given by $\eta_i(\boldsymbol{\theta}) = \theta_i$. If all $\theta_i > 0$, all species may coexist. Conversely, given only measurements of the equilibrium relative yields $\eta_i$, our best guess for interactions $\beta_{ij}$ is that they are drawn from $P(\boldsymbol{\beta}|\boldsymbol{\theta})$ with constraints (Eq. S5) and

$$\theta_i = \eta_i \qquad \text{(for Lotka-Volterra dynamics at equilibrium).} \tag{Eq. S6}$$

Following the mathematical derivation in Sec. B.4, we find that interactions $\beta_{ij}$ should follow two statistical patterns which both admit intuitive interpretations (Fig. S2). First, interactions are, on average, biased in a way that explains each species' success: given the relative yields of species $i$ and $j$, the expectation of $\beta_{ij}$ differs from the mean of the prior distribution, $\bar{\beta}$, as

$$\Delta\beta(\theta_i, \theta_j) = \mathrm{E}[\beta_{ij}|\theta_i, \theta_j] - \bar{\beta} = -\frac{(\theta_i - \theta^*)\theta_j}{\sum_{m\neq i}\theta_m^2}(1 - \bar{\beta}) \tag{Eq. S7}$$

Here, $\theta^* = (1 - \bar{\beta}\sum_i \theta_i)/(1 - \bar{\beta})$ is a threshold indicating which species are biased for or against in the competition matrix. When $\theta_i > \theta^*$, a species suffers less competition than the prior mean, and conversely if $\theta_i < \theta^*$. Importantly, we see that this bias is not evenly distributed: competition coming from large $\theta_j$ species tends to be more biased.

The interpretation of $\theta^*$ becomes more intuitive when we recall that, in the Lotka-Volterra model used here, each species $i$ will attain $\eta_i = \theta_i$ at equilibrium.

Therefore, in the main text, we use $\eta_i$ as estimates of $\theta_i$, and we have a threshold $\eta^*$ such that

$$0 = 1 - \eta^* - \bar{\beta}(\eta_{tot} - \eta^*) \quad \text{(Eq. S8)}$$

which means that $\eta^*$ is the relative yield that we would expect for a species, given the relative yield total $\eta_{tot} = \sum_i \eta_i$, if all the interactions of that species with others were exactly equal to the prior mean $\bar{\beta}$. Having $\eta_i > \eta^*$ then intuitively suggests that interactions suffered by $i$ should on average be less competitive (note however that this is not a mathematical necessity, only the most intuitive and parsimonious possibility).

The second pattern imposes that successful species $j$ and $k$ avoid competing against the same species $i$,

$$\mathrm{corr}(\beta_{ij}, \beta_{ik}|\theta_i, \theta_j, \theta_k) = -\frac{\theta_j \theta_k}{\sum_{m \neq i} \theta_m^2}. \quad \text{(Eq. S9)}$$

The first pattern (Eq. S7) decides the *expected* success of each species, this second equation guarantees that the equilibrium value is *exactly* set to $\eta_i(\boldsymbol{\theta}) = \theta_i$, as explained in Fig. S3 and in Sec. B.4.

#### B.3 Discussion

##### B.3.1 How to interpret the pattern

Since there is a single parameter $\theta_i$ associated with each species (by contrast with coexistence mechanisms that have more parameters than species, and parameters that are not linked to any one species), we call $\theta_i$ the "position" of species $i$ in the network, and label it as a "core" position if the species participates strongly in the diffuse clique (high $\theta$) and a "satellite" position if it does not.

Naively, one might assume that coexistence requires all competitive interactions to be much weaker than in random assemblages of non-coexisting species. But finding a set of nothing but weak competitors is unlikely if migration allows others to enter the community. Instead, the most likely and parsimonious outcome is the differentiation of species into core and satellite (high and low $\theta$), where the former are both more discriminate in their competition and more successful at equilibrium. Rather than have all species deviate strongly from the global distribution, this deviation is thus borne mostly by some species (those with high $\theta$), in a way that ensures that they are dominant. This concentrates the non-random constraints into a fraction of important species, while allowing maximal random variation around this structure.

This is the most parsimonious or least constraining arrangement of interactions that allows all species to cohabit. Therefore, this pattern is the one that we find in most possible solutions to the problem of coexistence (Fig. 1 in the main text). When we simulate ecological dynamics starting from a random pool of species, we see the same pattern emerge among those species that can survive together (Fig. S4). We interpret this pattern to mean that core species are more distinctly partitioned into ecological niches – at least, phenomenological niches arising from many latent factors, if not necessarily identifiable biological mechanisms – whereas satellite species interact in a more diffuse fashion (Fig. S4b). However we stress that $\theta_i$ is not a trait of the species itself, but a property of its embedding in an interaction network, and the same species will under all likelihood be associated with different values of $\theta_i$ in different communities.

##### B.3.2 Why the pattern emerges

The coexistence of many competitors at equilibrium can result from a vast range of interaction matrices $\boldsymbol{\beta}$, reflecting all sorts of underlying trade-offs. Requiring that all species survive imposes $S$ constraints, whereas there are $S(S-1)$ interaction coefficients $\beta_{ij}$. Hence, many different sets

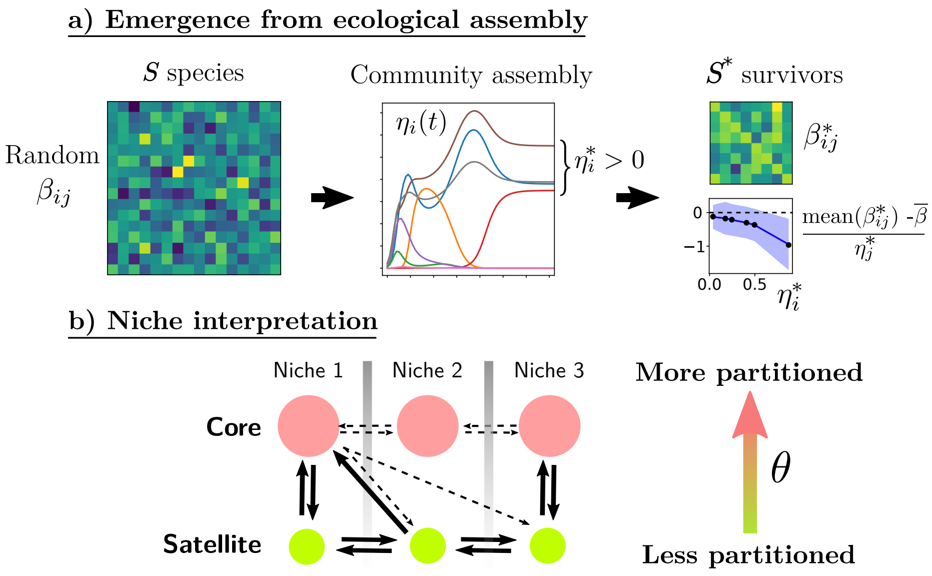

Figure S4: Interpreting diffuse partitioning. **a)** Emergence of the pattern from ecological dynamics. Running the dynamics (Eq. S2) from a pool of $S$ species with random interactions, we find that the interaction matrix $\boldsymbol{\beta}^*$ restricted to the $S^*$ survivors exhibits the expected statistical relations (Eq. S7) and (Eq. S9). **b)** This pattern can be interpreted as reflecting an underlying niche structure, although these phenomenological niches likely emerge from the interplay of many factors, and may not be identifiable to particular biological mechanisms. Core (high $\theta$) species then appear more strongly partitioned, in the sense that they compete weakly across niche boundaries, while satellite species compete more randomly against all species. Note that position $\theta_i$ is not an intrinsic trait of species $i$, and is expected to vary for the same species in different communities.

of coefficients can produce the same results, although they may correspond to distinct mechanisms. The probabilistic argument above succeeds because, in this large ensemble of possible matrices $\boldsymbol{\beta}$, most coefficients $\beta_{ij}$ appear to be close to randomness (minimally biased by the constraints) rather than ordered in particular ways [**?**].

That is not yet enough to explain why our statistical pattern should hold within one specific community. We must invoke the further property of *self-averaging*, which is key to simplicity in high-dimensional systems: as the size $S$ of the community increases, statistics within one community become closer to statistics over all communities.

#### B.4 Mathematical derivation of the pattern

**Outline:** To sample $M$ random numbers that satisfy $N$ constraints, we can draw $M-N$ independent random numbers, then translate and rotate them to occupy a $M$-dimensional space. For instance, the two random numbers $b_1$ and $b_2$ in Fig. S3 could have been drawn as a single random number $b$ along a line, then translated and rotated to have two values that always respect the constraint $b_1+b_2-1=0$.

The probability distribution of coefficients in the matrix $\beta$ has $\beta_{ii}=1$, with priors $\langle\beta\rangle \equiv \langle\beta_{ij}\rangle$ and $\left\langle\beta^2\right\rangle_c \equiv \left\langle\beta_{ij}^2\right\rangle_c$, and conditioned on

$$\sum_{j,(j\neq i)} \beta_{ij}\eta_j + \eta_i = 1$$

for every $i$. Consider $\vec{\beta}_i$, the $i$-th row without $\beta_{ii}$ (it has $S-1$ elements). The probability distribution for $\gamma=0$ depends on $\eta_i$ and the vector of abundances without $\eta_i$, denoted $\vec{\eta}_{-i}$, and given by

$$P\left(\vec{\beta}_i|\vec{\eta}\right) \propto \text{Normal}\left(\vec{\beta}_i; \left\langle\beta^2\right\rangle_c \mathbf{I}, \vec{\mu} = \langle\beta\rangle\,\vec{u}\right)\ \delta\left(\vec{\beta}_i\cdot\vec{\eta}_{-i} - (1-\eta_i)\right)$$

where $\vec{u}$ is the column vector $\vec{u}=(1,1,...,1)^T$, $\delta$ is the Dirac delta function, and we do not make explicit the normalization factor which doesn't depend on the $\beta_{ij}$-variables. We would like to write this as a Gaussian probability distribution without a delta-function. Instead, it will have a rank-deficient correlation matrix $\mathbf{C}$.

To proceed, consider an orthonormal change of basis:

$$\vec{x} = \mathbf{R}\vec{\beta}_i\ .$$

We choose the first row of $\mathbf{R}$ to be $\vec{a} \equiv \vec{\eta}_{-i}/\left|\vec{\eta}_{-i}\right|$, and the rest of the rows are chosen orthogonal to it and normalized (e.g., via a Gram-Schmidt process). Thus $\mathbf{R}\vec{\eta}_{-i} = \left|\vec{\eta}_{-i}\right|\vec{w}$, with $\vec{w}\equiv(1,0,...,0)^T$. In the rotated space,

$$\vec{\beta}_i\cdot\vec{\eta}_{-i} = \left(\mathbf{R}\vec{\beta}_i\right)\cdot\left(\mathbf{R}\vec{\eta}_{-i}\right) = \left|\vec{\eta}_{-i}\right|\vec{x}\cdot\vec{w} = \left|\vec{\eta}_{-i}\right| x_1\ .$$

Therefore, the distribution of $\vec{x}$ is:

$$P(\vec{x}) = \text{Normal}\left(\vec{x}; \left\langle\beta^2\right\rangle_c \mathbf{I}, \mathbf{R}\langle\beta\rangle\,\vec{u}\right)\ \delta\left(\left|\vec{\eta}_{-i}\right| x_1 - (1-\eta_i)\right)\ .$$

We can readily rewrite this as a Gaussian distribution without the delta-function, since it represents $S$ independent random variables with a constraint only on the first. The loss of variance in the first dimension means that the correlation matrix becomes

$$\mathbf{C}_x = \left\langle\beta^2\right\rangle_c \mathbf{I} - \left\langle\beta^2\right\rangle_c \begin{pmatrix} 1 & & & \\ & 0 & & \\ & & 0 & \\ & & & \ddots \end{pmatrix}\ .$$

While in the means vector, $\vec{\mu}_x$, we have $\langle x_1 \rangle = (1-\eta_i) / \left|\vec{\eta}_{\setminus i}\right|$ and the rest are unchanged,

$$\vec{\mu}_x = \left[\mathbf{I} - \begin{pmatrix} 1 & & & \\ & 0 & & \\ & & 0 & \\ & & & \ddots \end{pmatrix}\right] \mathbf{R} \langle \beta \rangle \vec{u} + \vec{w}\,(1-\eta_i) / |\vec{\eta}_{-i}| \ .$$

Together,

$$P(\vec{x}) = \text{Normal}\left(\vec{x}; \mathbf{C}_x, \vec{\mu}_x\right) \ .$$

Rotating back to $\vec{\beta}_i = \mathbf{R}^{-1}\vec{x} = \mathbf{R}^T\vec{x}$ we find

$$P\left(\vec{\beta}_i | \vec{\eta}_{-i}\right) = \text{Normal}\left(\vec{\beta}_i; \mathbf{C}_{\vec{\beta}_i}, \vec{\mu}_{\vec{\beta}_i}\right) \ ,$$

with

$$\mathbf{C}_{\vec{\beta}_i} = \mathbf{R}^T \mathbf{C}_x \mathbf{R} = \left\langle \beta^2 \right\rangle_c \left[\mathbf{I} - \vec{a}\vec{a}^T\right] = \left\langle \beta^2 \right\rangle_c \left[\mathbf{I} - \frac{\vec{\eta}_{-i}\vec{\eta}_{-i}^T}{|\vec{\eta}_{-i}|^2}\right] \ ,$$

$$\vec{\mu}_{\vec{\beta}_i} = \mathbf{R}^T \left\{ \left[\mathbf{I} - \begin{pmatrix} 1 & & & \\ & 0 & & \\ & & 0 & \\ & & & \ddots \end{pmatrix}\right] \mathbf{R} \langle \beta \rangle \vec{u} + \vec{w} \frac{(1-\eta_i)}{\left|\vec{\eta}_{\setminus i}\right|} \right\}$$

$$= \left[\mathbf{I} - \vec{a}\vec{a}^T\right] \langle \beta \rangle \vec{u} + \vec{a} \frac{(1-\eta_i)}{|\vec{\eta}_{-i}|} = \langle \beta \rangle \left[\vec{u} - \frac{\vec{\eta}_{-i}\vec{\eta}_{-i}^T}{|\vec{\eta}_{-i}|^2}\vec{u}\right] + \frac{(1-\eta_i)}{|\vec{\eta}_{-i}|^2}\vec{\eta}_{-i}$$

Written element-wise they read

$$\mathbf{C}_{\vec{\beta}_i}(k,l) = \left\langle \beta^2 \right\rangle_c \left[\delta_{k,l} - \frac{\eta_k \eta_l}{\sum_{j,(j\neq i)} \eta_j^2}\right]$$

$$\vec{\mu}_{\vec{\beta}_i}(k) = \langle \beta \rangle - \langle \beta \rangle \frac{\eta_k \sum_{l,(l\neq i)} \eta_l}{\sum_{j,(j\neq i)} \eta_j^2} + \frac{(1-\eta_i)}{\sum_{j,(j\neq i)} \eta_j^2}\eta_k$$

$$= \langle \beta \rangle + \frac{1 - \eta_i - \langle \beta \rangle \sum_{l,(l\neq i)} \eta_l}{\sum_{j,(j\neq i)} \eta_j^2}\eta_k$$

$$= \langle \beta \rangle - \frac{1}{\sum_{j(\neq i)}^S \eta_j^2}\eta_i\eta_k + \frac{1 - \langle \beta \rangle \sum_{j(\neq i)}^S \eta_j}{\sum_{j(\neq i)}^S \eta_j^2}\eta_k$$

which is precisely what we found above for the change in means. But now we also have the correlations between the elements, which also agree with assembly for large $S$. The correlations can be easily intepreted as the projector that kills fluctuations in the $\sum_j \beta_{ij}\eta_j$ direction.

The derivation can be turned into an algorithm: the matrix $\mathbf{R}$ can be generated and $\vec{x}$ can be easily sampled, as implemented in R in the code that we provide in main text.

#### B.5 Extension for interaction symmetry

All expressions in the main text were given under the assumption of asymmetrical interactions, $\gamma = \text{corr}(\beta_{ij}, \beta_{ji}) = 0$ (which is borne out in our analysis of the empirical data). The general expression derived in [?] is simply a symmetrization of the prediction for $\gamma = 0$

$$E[\beta_{ij}|\theta_i,\theta_j,\gamma] = \overline{\beta} + \left(E[\beta_{ij}|\theta_i,\theta_j,\gamma=0] - \overline{\beta}\right) + \gamma\left(E[\beta_{ji}|\theta_i,\theta_j,\gamma=0] - \overline{\beta}\right) \qquad \text{(Eq. S10)}$$

This symmetrization induces correlations

$$\text{corr}(\beta_{ij}, \beta_{ik}|\theta_j, \theta_k) = \delta_{jk} - A_i\theta_j\theta_k \tag{Eq. S11}$$

$$\text{corr}(\beta_{ij}, \beta_{ki}|\theta_j, \theta_k) = -\gamma A_i\theta_j\theta_k \tag{Eq. S12}$$

$$\text{corr}(\beta_{ji}, \beta_{ki}|\theta_j, \theta_k) = -\gamma^2 A_i\theta_j\theta_k \tag{Eq. S13}$$

with $A_i = 1/\sum_{j\neq i}\theta_j^2$. We can thus extend the method to empirical settings where interactions would be significantly symmetrical or antisymmetrical, which is not the case here.

#### B.6 Generating the competition-colonization tradeoff matrices

In the main text, we compare the high-dimensional diffuse clique structure to the one-dimensional competition-colonization tradeoff mechanism [**?**, **?**]. In this tradeoff, each species is characterized by two parameters, $c_i$ and $m_i$ representing its competitive rank and mortality. These parameters determine the interaction matrix used in our work:

$$\beta_{ij} = \frac{(c_i + c_j)(c_j - m_j)}{(c_i - m_i)c_j} \tag{Eq. S14}$$

The values of $c_i$ and $m_i$ for each species $i$ must follow a precise set of inequalities to ensure the coexistence of all species [**?**].

An interaction matrix for $S$ species respecting these inequalities can be generated iteratively as follows:

1. We select scales $m = 1$ and $c = 2$ for mortality and competition

2. We initialize a sequence $x = \{x_i\}$ with $x_1 = 1$

3. For each species $i$ starting from $i = 1$, its competitive ability is computed as

$$c_i = 2 + \frac{m}{\left(1 - \frac{c-m}{c}\sum_{j\leq i} x_j\right)^2} \tag{Eq. S15}$$

4. We then set the competition coefficients to

$$\beta_{ij} = \frac{(c_i + c)(c - m)}{c(c_i - m)} \tag{Eq. S16}$$

   for all $j < i$, $\beta_{ii} = 1$ and $\beta_{ij} = 0$ for all $j > i$.

5. Finally we add a new element to the sequence $x$,

$$x_{i+1} = 1 - \sum_{j\leq i} \beta_{ij} x_j \tag{Eq. S17}$$

   and iterate until all species have been considered.

### C Description of the experiments

The designs of the three biodiversity experiments used are summarized in Table S1 for ease of comparison.

| | Big Bio | BioCON | Wageningen |
|---|---|---|---|
| Year established | 1994–1995 | 1998 | 2000 |
| Years used | 1997–2015 | 2000-2017 | 2002–2011 |
| Establishment | seeds | seeds | transplanted seedlings |
| Diversity levels | 1,2,4,8,16 | 1,2,4,9,16 | 1,2,4,8 |
| Number of species | 18 | 16 | 8 |
| Legumes present | yes | yes | no |
| Number of plots | 168 | 359 | 102 |
| Monoculture replication | 1–3 depending on species | 2 | 6 |
| Plot size | 9 x 9 m | 2 x 2 m | 1 x 1 m |
| Samples per plot | 4 in 2001–2006 $^{n1}$ | 1 | 1 |
| Sample size | Size of clipped strips have periodically changed | 0.1 x 1 m | 0.6 x 0.6 m |

Table S1: Summary of the experimental designs of the three long-term grassland biodiversity experiments.

#### C.1 Wageningen experiment

The experiment near Wageningen, The Netherlands was established during 2000 [?, ?]. In each plot, the topsoil was removed to a depth of 50 cm. Wooden frames were placed around the edges of these holes, which were 1 x 1 x 0.5 m (length x width x height), and each hole was filled with a mixture of pure sand and soil (3:1) from an old field. Seedlings were grown in a greenhouse, and 144 seedlings were transplanted into each 1 x 1 m plot in a substitutive design (i.e., same density in all plots). The experiment consisted of 102 plots, planted in 6 blocks. Each block includes each of the 8 study species in monoculture, four 2-species mixtures, four 4-species mixtures, and the species mixture of all 8 species. No legumes were included in this study. Species composition was maintained by hand weeding. Our analysis includes aboveground biomass collected from 2000-2011. At each harvest, the 0.6 x 0.6 m interior of each plot was sampled. Previous studies have documented a drought in 2006 [?], which drove us to consider whether to remove such extreme years in our analysis in the following sections.

#### C.2 Cedar Creek experiments

##### C.2.1 BigBio

The main biodiversity experiment (`http://www.cedarcreek.umn.edu/research/data/methods?e120`) at the Cedar Creek Ecosystem Science Reserve, Minnesota, USA, was established in 1994-1995 [?]. Land was treated with herbicide, burned, bulldozed, ploughed and harrowed in 1993 to clear extant plants and minimize the accumulated seed bank. Each of the 168 plots (9 x 9 m) was seeded in 1994 with 10g seed/$m^2$ and in 1995 with 5g seed/$m^2$, with this mass divided evenly between species randomly selected from an 18-species pool. Plots were burned annually, and included 1, 2, 4, 8, or 16 species. Species composition was maintained by hand weeding. Measurements used in our analysis were measurements of total aboveground biomass, collected all years, and species-specific biomasses, collected from 2000-2010. At each harvest, aboveground biomass was measured in 4 samples within each plot. During 1996-1999, each sample was 0.1 x 3 m; during 2000-2010, each sample was 0.1 x 6 m.

There were four complications in applying our methodology to these data. First, there were three species with non-replicated monocultures due to lack of seed establishment: *Elymus canadensis*, *Poa pratensis* and *Panicum virgatum*. Secondly, oaks were excluded from our analysis (*Quercus ellipsoidalis* and *Quercus macrocarpa*) since their growth patterns differed qualitatively from that of the other vegetation, and two other species (*Elymus canadensis* and *Agropyron smithii*) were excluded because their monocultures became dominated by other species. This was accomplished by excluding all plots that were dominated by these species ($> 50\, g/m^2$ average biomass for these species during 2000-2010) and ignoring their biomasses elsewhere. Third, due to seed contamination the legumes *P. villosum* and *P. candidum* were both sown in approximately equal densities in the 2-, 4-, and 8-species mixtures. Also, *Amorpha canescens* was sown instead of *P. villosum* in 16-species mixtures. Our analysis treats the biomass of these species as a single compound species. Fourth, due to the non-legume forb *Solidago rigida* not germinating in 1994, plots containing this plant were seeded with the non-legume forb *Monarda fistulosa* in 1995. *S. rigida* germinated the following year, hence all plots originally intended to be planted with *S. rigida* contained both species. Our analysis treats the biomass of these species as a single compound species.

With these provisions, there remained 24 distinct 16-species mixtures (out of 35 combinations in the data and 153 possibilities for a 18-species pool). We treated each mixture as a distinct test set (Set B in Sec. A.3), against which we tested the same inferred interaction coefficients $\beta_{ij}$ obtained from plots with $S < 16$. The agreement metrics shown in main text represent the distribution over all 16-species mixtures.

##### C.2.2 BioCON

The BioCON (Biodiversity, $CO_2$ and N) experiment (`http://www.biocon.umn.edu/`) is located at the Cedar Creek Natural History area in Minnesota, USA. Plots were established in 1997 on a secondary successional grassland. The experimental area was tilled and fumigated with methyl bromide. The experimental treatments were arranged in complete factorial combination of $CO_2$ (ambient or 560mmol mol$^{-1}$), species number (1, 4, 9, and 16, chosen from a pool of 16 species) and N level (control and fertilized). Each plot was seeded in 1997 with 12 $gm^{-2}$ of seed partitioned equally among all species assigned to a plot.

Treatments are arranged in a split-plot design. $CO_2$ treatment is the whole-plot factor and is replicated three times among the six rings. The subplot factors of species number and N treatment were assigned randomly and replicated in individual plots among the six rings. For each of the four combinations of $CO_2$ and N levels, pooled across all rings, there were 32 randomly assigned replicates for the plots planted to 1 species, 15 for those planted to 4 species, 15 for 9 species, and 12 for 16 species. An additional functional diversity experiment had another 63 plots with 4 species, resulting in a total of 359 plots used for our analysis.

Beginning in 1998, N addition plots received 4 $g_m^{-2}yr^{-1}$ N three times per year. $CO_2$ was added using FACE technology. Some of the 9 species plots received water shading treatments starting 2007 and or warming treatments starting 2012 (http://www.cedarcreek.umn.edu/research/data/methods?e141); these treatments were excluded from our analysis. Above-ground biomass was harvested by clipping a 10 × 100 cm strip just above the soil surface and sorting to species. This was measured in June and August from 1998 to 2011, and in August from 2012 to 2017.

For our analysis, we only considered the biomasses measured in August. We removed species whose biomass was reported as 0 in more than 95% of the cases (plots and years) where they had been seeded: *Anemone cylindrica* and *Petalostemum villosum* for all treatements, and *Amorpha canescence* for the treatment "+C+N". The four treatments were analyzed separately and are designated as "biocon", "biocon+C", "biocon+N" and "biocon+C+N" in Fig. S7 and **??**. The agreement metrics shown in main text represent the distribution over all treatments, while scores per treatment are shown in Appendix F below.

#### C.3 Estimating equilibrium abundances

Our results require estimating the abundances at equilibrium for each species in each composition in the dataset. However, there are many sources of both measurement errors and dynamical fluctuations in the experiments, hence we need some scheme to evaluate the underlying equilibrium abundances.

After investigating many possibilities, we settled (as described in Sec. A.3) on a very simple scheme: estimating equilibrium abundance $N_i(w)$ in plots with composition $w$ as the arithmetic mean over all abundance values observed for species $i$ at different times and in different plots with the same composition. The only additional subtleties are that we remove the first 2 years in all experiences in order to bypass transient dynamics linked to species establishment, and we use bootstrapping to get multiple estimates of the mean.

To explain this final choice of a simple scheme, we detail here the different possibilities that we have considered in how to bin, filter and pre-process the data before taking the mean and error on the mean as estimates of equilibrium abundance:

1. Keeping one vector per replica plot while filtering out-of-equilibrium years.
   We systematically tested the following possibilities:

   (a) Drop the first two years

   (b) Drop extreme years (e.g. the drought year 2006 in the Wageningen experiment [?])

   (c) Retain all years

2. Making one vector from all the replicas, then fitting a functional form, and computing variance around the trend.

   - Two possible functional forms: saturating expression (solution of logistic) and exponential decay, where $t = y - y_0$, $y_0$ being the first year

   $$x(t) = \frac{a}{1 - be^{-ct}}, \qquad x(t) = a + be^{-ct} \tag{Eq. S18}$$

   - Distance-based selection between logistic and exponential decay to $a = 0$ to detect extinctions. (Fig. S5)

3. One vector per replica plot, but detrended to remove the effect of global fluctuations of biomass:

   - Removing the trend in carrying capacities (**detrendK** in Fig. S6), by dividing the abundance of species $i$ at any time $t$ and in any plot by the median of its monoculture abundances at the same time $t$
   - Removing the trend in total biomasses (**detrendall** in Fig. S6), by dividing the abundance of species $i$ at any time $t$ and in any plot by the median of all biomasses of all species in all plots at the same time $t$
   - Removing the trend in growth rates (**detrendret** in Fig. S6), by dividing the abundance of species $i$ at any time $t$ and in any plot by a constant such that the distribution of growth of all species in all plots between times $t - 1$ and $t$ has a median of 0

In the end, a battery of tests (a few of which are illustrated in Fig. S6 and detailed in Sec. D.3) convinced us that all these different preprocessing techniques either did not improve the quality of our estimate of equilibrium abundances, or made it worse – with the notable exception of removing the first two years, as they are clearly out of equilibrium in most experiments (see Figs. S5 and S6 for Wageningen), and choosing the same procedure for all experiments was intended to limit the bias introduced between experiments.

### D Fitting the interactions

#### D.1 Lotka-Volterra model

The standard competitive Lotka-Volterra model takes the form [?]

$$\frac{1}{N_i}\frac{dN_i}{dt} = r_i \left(1 - \frac{N_i - \sum_{j \neq i} \alpha_{ij} N_j}{K_i}\right) \tag{Eq. S19}$$

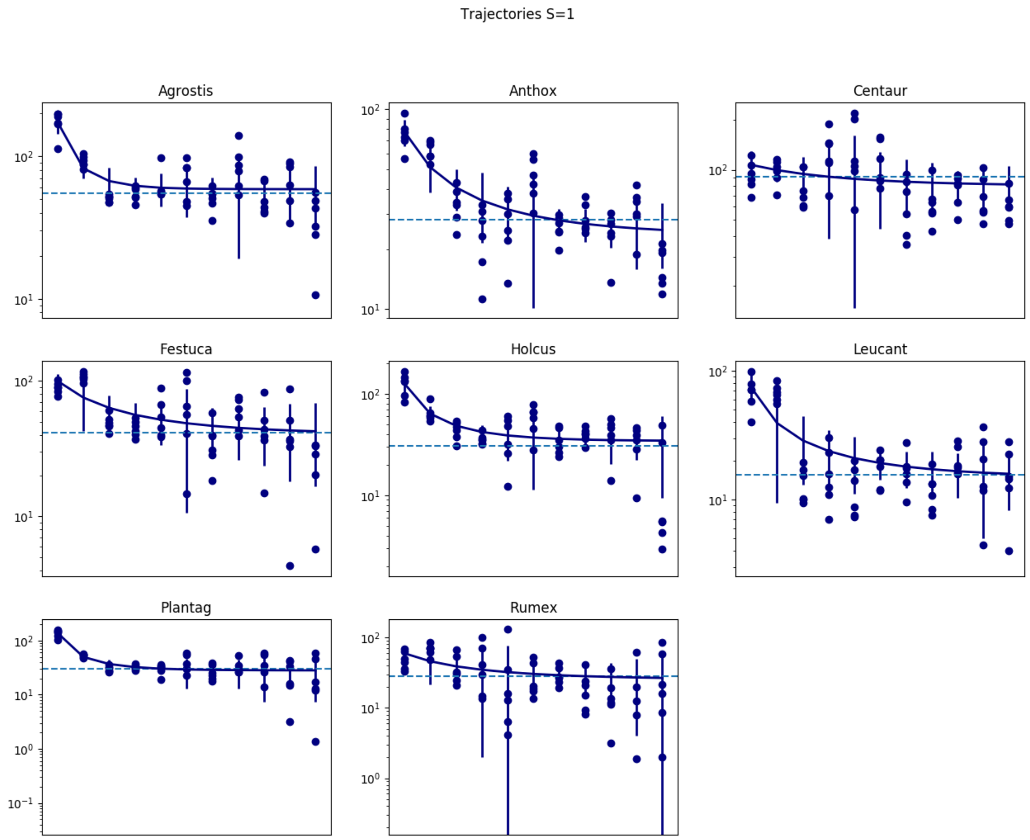

Figure S5: Trajectories in monocultures in the Wageningen grassland experiment and example of fit. We can see that the first two years at least are distinctly out of equilibrium for many trajectories (this is also true for polycultures). In addition, we show here the good agremeent between the carrying capacity estimated as the asymptotic value of the fitted monoculture trajectory, and the value inferred using all plots in Sec. D (dashed lines).

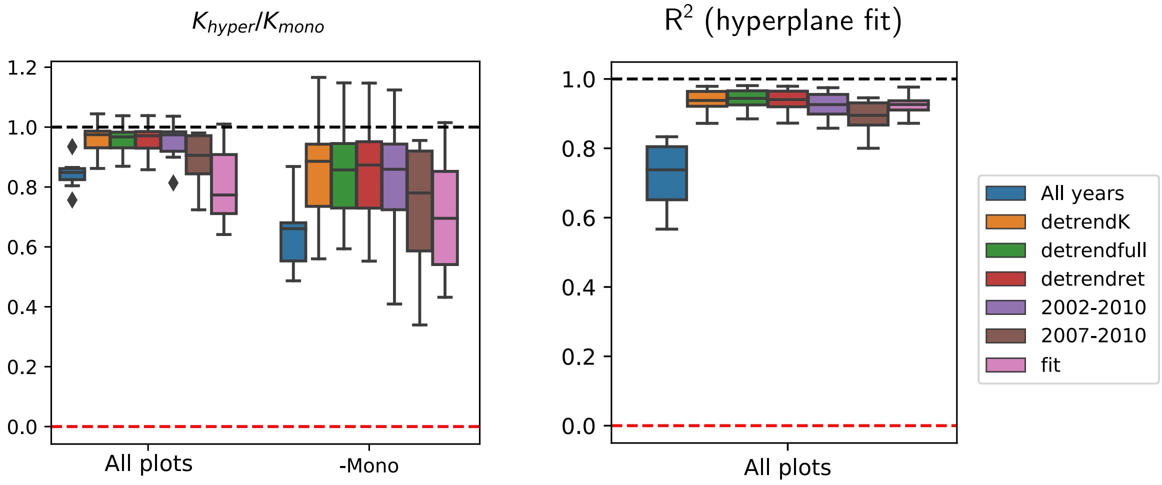

Figure S6: Indicators of success of hyperplane regression (Sec. D.2), depending on how Wageningen experiment data was processed to estimate the equilibrium abundance of each species in each plot. **Left panel:** We show a bar plot the ratio of $K_{hyper}$ (carrying capacities inferred as the intercept of the hyperplane fit, see Sec. D.2) and $K_{mono}$ (carrying capacities measured in monocultures), when the hyperplane inference includes all plots (left) and without monocultures (right). **Right panel:** We show the $R^2$ of the hyperplane fit. Processing includes dropping some years and detrending (see Sec. C.3). From both panels, we reach a common conclusion: all choices of how to process the data are close to being equivalent for the Wageningen experiment, except taking averages over all years (because of the out-of-equilibrium first two years, see Fig. S5), and estimating equilibrium abundances from fits of the trajectories (these fits are successful for monocultures. The agreement also decreases somewhat if we remove more than 2 years (for instance taking only data points after the 2006 drought).

In a setting with a wide distribution of carrying capacities $K_i$, the previous expression will allow a very low fraction of survivors if the $\alpha_{ij}$ are distributed from some narrow (e.g. normal) distribution [?]. Thus, it is often more meaningful to consider the abundance relative to the carrying capacity $\eta_i = N_i/K_i$ (also known as $r$elative yield), for which the equation becomes

$$\frac{1}{\eta_i}\frac{d\eta_i}{dt} = r_i\left(1 - \sum_j \beta_{ij}\eta_j\right) \qquad \text{(Eq. S20)}$$

where $\beta_{ij} = \alpha_{ij}K_i/K_j$ are rescaled interactions, and by definition $\beta_{ii} = 1$. Since we do see rare and abundant species coexist stably, we expect $\eta_i$ and $\beta_{ij}$ to be better variables than $N_i$ and $\alpha_{ij}$ to understand the dynamics, see also SI in [?].

#### D.2 Hyperplane method

Given the biomasses $N_i^{(1)}$ in monocultures and $N_i^{(w)}$ in other plots, we define the relative yield

$$\eta_i^{(w)} = N_i^{(w)}/N_i^{(1)}. \qquad \text{(Eq. S21)}$$

In principle there should be a single such value per composition, since we will take it to represent the true equilibrium relative yield of that species in that composition. In practice, as detailed in Sec. A.3, we obtain error bars on the $\boldsymbol{\eta}$ and $\boldsymbol{\beta}$ by bootstrapping: we generate many different vectors of abundances $N_i^{(w)}$ and $N_i^{(1)}$ by resampling with replacement from the set of values $N_i^{(w)}$ for different years and replica plots (vectors generated this way have the same number of entries as the original data), then take the mean of each vector, and take their ratio to compute one value of $\eta_i^{(w)}$. The bootstrap process creates many such vectors, and therefore many values of $\eta_i^{(w)}$, and their variations are assumed to represent the standard error on the mean. The different generated values of $\eta_i^{(w)}$ are then used as follows to compute different matrices $\boldsymbol{\beta}$, and hence to obtain error bars on the coefficients $\beta_{ij}$.

In principle, we can deduce interactions from the relative yields in duoculture plots ($w$ such that $S_w = 2$)

$$\beta_{ij} = \frac{1 - \eta_i^{(w)}}{\eta_j^{(w)}}. \qquad \text{(Eq. S22)}$$

In practice, using only duoculture relative yields leads to rather noisy results.

Following Xiao et al [?] we instead infer the interaction matrix $\beta_{ij}$ as follows. Provided that species $i$ is not extinct, all equilibria containing that species are predicted (under Lotka-Volterra assumptions) to verify

$$0 = 1 - \eta_i^{(w)} - \sum_{j\neq i} \beta_{ij}\eta_j^{(w)} \qquad \text{(Eq. S23)}$$

where $w$ is the index of some plot with $S(w)$ species. Under the simplifying assumption of asymmetrical interactions, i.e. $\beta_{ij}$ independent of $\beta_{ji}$, we can thus infer the vector $\beta_{ij}$ for each species $i$ independently. It is then a simple case of multilinear regression, minimizing the sum of squares over all plots $w$

$$\min_{\beta_{ij}}\left(\sum_w \left(1 - \eta_i^{(w)} - \sum_{j\neq i}\beta_{ij}\eta_j^{(w)}\right)^2\right) \qquad \text{(Eq. S24)}$$

In the case of the Wageningen grassland experiment, for each species $i$, there are 7 parameters $\beta_{ij}$ and at most 30 plots $w$ in which the species is present with significant abundance. For other experiments, there are fewer different compositions per species, meaning that there is a larger risk of overfitting some interactions; in some, there are pairs of species that never occur in the same composition, hence their interactions are left blank and ignored in the inference.

The same process can be applied directly to species biomasses

$$0 = K_i - N_i^{(w)} - \sum_{j \neq i} \beta_{ij} N_j^{(w)} \tag{Eq. S25}$$

so as to infer the carrying capacity $K_i$, not from monocultures only, but as the intercept of the multilinear regression of $N_i$ against other biomasses $N_j$ (i.e. the asymptotic value of $N_i$ in the absence of other species). These estimates of $K_i$ are used in Sec. D.3 to test the validity of the fitted linear model.

#### D.3 Testing the Lotka-Volterra equilibrium model

We perform a series of tests to demonstrate the qualitative or quantitative validity of our particular choice of model for interactions (Eq. S2). These tests are most successful for the Wageningen experiment, which we therefore propose as a reference experiment for any further investigation based on Lotka-Volterra competition models.

In particular, carrying capacities $K_i$ inferred from all multispecies plots ($S > 1$) show good agreement with measurements in monocultures ($S = 1$), see Fig. S6. Likewise, interactions $\beta_{ij}$ inferred from all plots with $S < S_{\max}$ are compatible with the equilibrium values of $\eta_i = B_i/K_i$ in the full community ($S = S_{\max}$), see Fig. S7. This strongly supports the simple linear model (Eq. S3). In addition, the Wageningen experiment is by far the least sensitive to the choice of process used to estimate abundances and interactions. By contrast, data from other experiments (not shown) appear less robust and may be affected by nonlinearity, transient dynamics, stochasticity, and measurement errors.

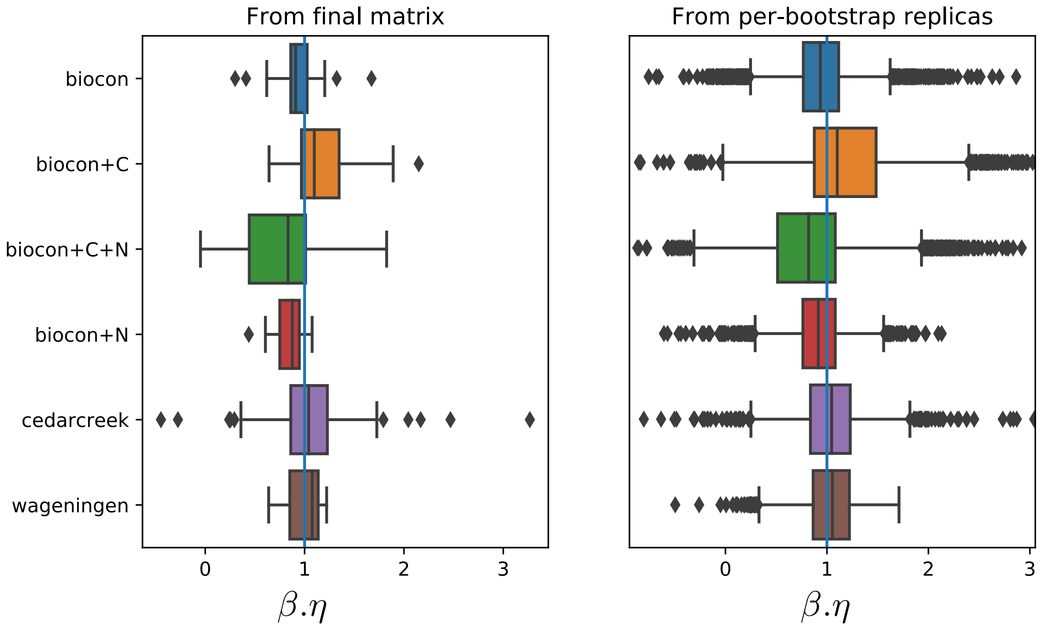

Figure S7: Distribution of $\sum_j \beta_{ij}.\eta_j$ for all species $i$ in each experiment. $\beta_{ij}$ is inferred from all plots with $S < S_{\max}$ (with $\beta_{ii} = 1$), while $\eta_i$ is measured in plots with $S = S_{\max}$. If $\sum_j \beta_{ij}\eta_j = 1$ for all $i$, then this means that the interactions inferred from subcommunities correctly predict the relative yields in the full community, i.e. the model of linear interactions at equilibrium is a valid description of the community. We see here that the Wageningen experiment is closest to always exhibiting $\beta.\eta \approx 1$ (values are both centered around 1 and narrowly distributed). The left panel shows results for all rows $i$ in the final interaction matrix inferred through our process (taking the median over all the coefficients $\beta_{ij}$ inferred in different bootstrapped replicates), while the right panel shows $\beta.\eta$ computed in all bootstrapped replicas separately, then pooled together. We see that the bootstrapping and aggregation procedure leads to very significant noise reduction compared to considering a single point estimate for each $\beta_{ij}$ (i.e. individiual replica).

### E Computing and testing theoretical predictions

We have two sets of predictions to test:

- a single-interaction pattern with $E[\beta_{ij}]$
- a two-interaction pattern with $\mathrm{corr}(\beta_{ij}, \beta_{ik})$

To test the first relationship, we plot the measured values of $y = \beta_{ij}$ against their predicted expectation $x = \overline{\beta} - (\eta_i - \eta^*)\eta_j / \sum_{k \neq i} \eta_k^2$. To obtain the empirical expectation, we perform a running average: we replace, for each point $(x, y)$, its $y$ by the average $\overline{y}$ within a window centered on $x$ and spanning 10% of the axis. We then group all values $\overline{y}$ associated with the same species pair $(i, j)$, take their median, and reconstruct an empirical matrix of expectations $B_{ij}$.

We proceed similarly to test the second relationship. Defining $d_{ij} = \beta_{ij} - E[\beta_{ij}|\eta_i, \eta_j]$, we compute $y = \delta_{jk} - d_{ij}d_{ik}/\mathrm{mean}\left(d_{mn}^2\right)$, where $\delta_{jk} = 1$ if $j = k$ and 0 otherwise, and the denominator is the sample mean. We then plot $y$ against the prediction $x = -\eta_j\eta_k / \sum_{l \neq i} \eta_l^2$, perform a running average to get $\overline{y}$, and construct an empirical tensor of correlations $C_{ijk}$ from the median of all values associated with each species triplet $(i, j, k)$.

The agreement is then estimated by looking at a correlative measure between empirical averages and theoretical expectations for each of the two relaitonships, as explained in Sec. E.4.

#### E.1 The why and how of bootstrapping

As described in Sec. A.3, we opted for a bootstrapping approach in order to account for inference error. The reason is that:

- theoretical predictions $E[\beta_{ij}|\eta_i, \eta_j]$ and $\mathrm{corr}(\beta_{ij}, \beta_{ik}|\eta_i, \eta_j, \eta_k)$ have some fluctuations that comes from errors in inferring the true equilibrium relative yields $\eta$
- measured coefficients $\beta_{ij}$ and $y_{ijk}$ have similar errors, also coming from inferred $\eta$, but they also exhibit true population variance: their exact error-free values would still be widely dispersed around the theoretical prediction

In consequence, when doing a plot, regression or other treatment of measured coeffiicents versus theoretical expectation, we cannot treat the two axes equally (a Type-2 regression is incorrect, for instance).

Instead, we choose to use bootstrapping to create many replicas of each point, with small displacements between these replicas due to the standard error in our inference. We then treat each replica as if it had zero error – in other words, as if the only variance was the true population variance, which we must average over only for measured coefficients (whereas theoretical expectations are treated as if they are known exactly for a given bootstrap replica).

#### E.2 Splitting the data to remove biases

For each bootstrapped replica of the data (see Sec. A.3) we split the data into two sets,

- **Set A:** used to obtain the empirical interaction coefficients $\beta_{ij}$ by regression combining many different compositions,
- **Set B:** used to predict the theoretical expectations $E[\beta ij]$ and $\mathrm{corr}(\beta_{ij}, \beta_{ik})$ only from species' relative yields in the full community

Since both methods rely on relative yields $\eta$, which require knowing carrying capacities $K$ (abundances in monoculture), we also split the monoculture data in two subsets: for each species, we pool together its observed abundances (at different dates and in different monoculture plots), then randomly distribute them into one half that will be used for set A, and another for set B.

This split is designed to try to reduce spurious correlations that could be induced by using the same monocultures to infer both the empirical $\beta_{ij}$ and the theoretical expectations. Yet this very split does introduce a weaker bias: since high values of $K$ will be found either in set **A** or in set **B** but not in both, we tend to create an anticorrelation between the median competition strength inferred in one and the other. Removing this second bias is done by randomly pairing set **A** from one bootstrapped replica to set **B** from another bootstrapped replica, to minimize both similarity and anti-similarity (avoidance) between the carrying capacities used in each part of our inference process.

#### E.3 Inferring prior mean

There are two important steps in computing the theoretical predictions: measuring the relative yields in the full community $\eta_i$, and inferring the parameter $\bar{\beta}$ which enters into our equation (Eq. S7). We now discuss the second step.

This parameter can be interpreted as the mean of the prior distribution of interactions $P(\boldsymbol{\beta})$ before we constrain it to ensure coexistence. It is therefore different from $E[\beta]$ the expected value of all interactions *among coexisting species* (see Sec. E.3.1). For instance, in a community assembly experiment where some species go to extinction, $\bar{\beta}$ would be the average interaction strength between all species in the pre-assembly pool, and $E[\beta]$ the average interaction strength in the observed community of surviving species, which would typically be less competitive.

Given the average interaction strength $E[\beta]$ between coexisting species, our theory [**?**] makes it possible to compute the original interaction strength $\bar{\beta}$ in the pool of all possible species (i.e. in the prior), see Sec. E.3.2.

We consider four methods for obtaining this parameter:

- Estimating $E[\beta]$ directly from the average of the empirical coefficients $\beta_{ij}$ (Sec. E.3.1), which is the option chosen in the main text. It is most direct, but requires knowledge of this interaction matrix, which is not available outside of special experimental setups such as those studied here[2]
- Estimating $E[\beta]$ only from the relative yields $\eta_i$ of the coexisting species in the maximum-diversity plots, by:
  - using a simple mean-field approximation (Sec. E.3.2)
  - assuming that the species we study constitute the full pool, so that $E[\beta] = \bar{\beta}$ (Sec. E.3.1)
  - simulating community assembly and selecting the prior parameters that lead to a distribution $P(\eta)$ closest to the observed one (Sec. E.3.3)

##### E.3.1 Change of mean and variance of $\beta$ due to coexistence

Let us rewrite the pattern of means (Eq. S7) as

$$E[\beta_{ij}|\eta_i, \eta_j] = \bar{\beta} + \frac{\eta_j}{S\left\langle \eta^2 \right\rangle - \eta_i^2}\left(1 - (1-\bar{\beta})\eta_i - \bar{\beta}S\left\langle \eta \right\rangle\right) \qquad \text{(Eq. S26)}$$

given the mean $\bar{\beta}$ of the prior distribution $P(\boldsymbol{\beta})$. We will compare this pattern to fits, and use it to generate matrices from the conditioned random ensemble. However, to do so, we must know $\bar{\beta}$.

[2]It also means that there is one element of information $E[\beta]$ transmitted between the Sets A and B as defined in Sec. A.3 before we compare their respective outcomes. In this case, to avoid creating biases, it is crucial to only transmit it once: we must take a single average over all the coefficients $\beta_{ij}$ obtained from all the bootstrapped replicas of Set A, then use that single number for all the predictions made using any bootstrapped replica of Set B. If we transmitted different estimates of $\bar{\beta}$ between replicas of Set A and replicas of Set B, we could 1) create correlations between pairs of replicas (which may be removed by randomization) and more importantly 2) artificially increase the variance of the theoretical predictions from set B.

By integrating over $\eta_i$ and $\eta_j$, we see that the average in the matrix of coexisting species is displaced from the prior mean

$$E[\beta_{ij}] = \bar{\beta}\left(1 + \langle\eta\rangle Q_2 - S\langle\eta\rangle^2 Q_1\right) + \langle\eta\rangle (Q_1 - Q_2) \qquad \text{(Eq. S27)}$$

with

$$Q_1 = \left\langle \frac{1}{S\langle\eta^2\rangle - \eta_i^2} \right\rangle, \qquad Q_2 = \left\langle \frac{\eta_i}{S\langle\eta^2\rangle - \eta_i^2} \right\rangle \qquad \text{(Eq. S28)}$$

As a consequence, by inverting (Eq. S27), we can infer $\bar{\beta}$ the mean in the prior distribution (or the pre-assembly pool of species) if we know the empirical average $E[\beta_{ij}]$ among coexisting species.

In particular, we can compute the *fixed point*: the value such that $E[\beta_{ij}] = \bar{\beta}$, meaning that adding the constraint of coexistence (or letting assembly run) will not change the average interaction. If we have no other information about the system, this is the most likely interaction strength given that we see these species coexist with observed relative yields $\eta_i$:

$$E[\beta_{ij}] = \bar{\beta} \quad \Leftrightarrow \quad \bar{\beta} = \frac{\langle\eta\rangle (Q_1 - Q_2)}{\langle\eta\rangle Q_2 - S\langle\eta\rangle^2 Q_1}. \qquad \text{(Eq. S29)}$$

This is one of our methods for estimating $\bar{\beta}$ and it gives good results in simulations and in the Wageningen experiment, but predicts weaker competition than computed directly from the matrix $\beta_{ij}$ in the other experiments, suggesting that species in these experiments may already represent a pre-selected subset of a larger, more competitive pool.

We also note that the correlation pattern

$$\mathrm{cov}(\beta_{ij}, \beta_{ik}) = \sigma_\beta^2 \left(\delta_{jk} - \frac{\eta_j \eta_k}{S\langle\eta^2\rangle - \eta_i^2}\right) \qquad \text{(Eq. S30)}$$

given the variance $\sigma_\beta^2$ of the prior distribution indicates that

$$\mathrm{var}(\beta) = \sigma_\beta^2 \left(1 - \langle\eta^2\rangle Q_1\right) \qquad \text{(Eq. S31)}$$

which we can invert to obtain $\sigma_\beta$ from a realized matrix.

##### E.3.2 Mean field estimate

Getting a naive estimate of $\bar{\beta}$ is immediate, following a classic *mean field approximation* [?]: if all interactions are exactly $\beta_{ij} = \bar{\beta}$ (i.e. if our prior distribution $P(\beta)$ has zero variance), then all relative yields should be equal and given by

$$\eta_i = \bar{\eta} = \frac{1}{1 + (S-1)\bar{\beta}} \qquad \text{(Eq. S32)}$$

We can invert this formula to obtain $\bar{\beta}$, but for heterogeneous relative yields, this parameter is strongly underestimated when compared to other methods (and also when compared to its true value in simulations). For a better (but still rough) approximation, we use the observed average relative yield $\langle\eta_i\rangle$ to estimate the *realized* interaction strength

$$E[\beta_{ij}]_{naive} = \frac{1}{S}\left(\frac{1}{\langle\eta_i\rangle} - 1\right) \qquad \text{(Eq. S33)}$$

From this, we can estimate $\bar{\beta}_{naive}$ by inverting (Eq. S27) as described in the previous section.

This estimate gives similar but slightly worse results than those shown in the main text, because it often underestimates the average strength of competition in a heterogeneous community. Note that if all species survive, we can also have a rough estimate of the variance of $\beta$ [?]:

$$\mathrm{var}(\beta)_{naive} = \frac{1}{S}\left(1 - \frac{\langle\eta\rangle^2}{\langle\eta^2\rangle}\right) \qquad \text{(Eq. S34)}$$

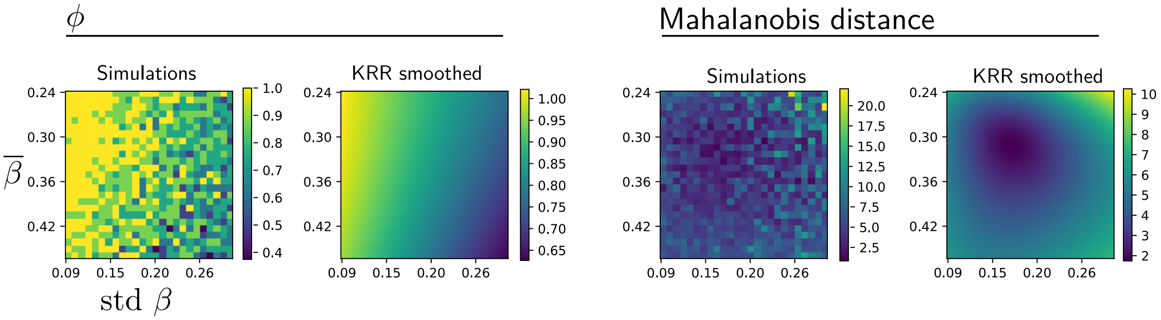

Figure S8: Measurements from simulations, and smoothed version obtained by Kernel Ridge Regression (implemented in Python with the scikit-learn library). We show $\phi$ (fraction of surviving species) and the Mahalanobis distance combining three statistics of $\eta_i$ including $\phi$, as defined in Sec. E.3, in the plane ($\bar{\beta}$,var($\beta$)) at $\gamma = 0$. The minimum of this distance gives the most likely value of $\bar{\beta}$.

##### E.3.3 Estimate from community assembly simulations

To obtain a better prior, we do the following (see Fig. S8) for each experimental dataset independently:

- We compute $\bar{\beta}_{naive}$ and $\text{var}(\beta)_{naive}$ from the above expressions
- We explore a region of the parameter space $(\bar{\beta}, \text{var}(\beta), \gamma)$ where $\gamma = \text{corr}(\beta_{ij}, \beta_{ji})$ is the interaction symmetry defined in Sec. B.5, around the initial values $\bar{\beta}_{naive}$, $\text{var}(\beta)_{naive}$ and $\gamma = 0$.
- at each point of this region, we run a simulation of a random Lotka-Volterra community, where interactions are drawn from a multivariate Gaussian distribution with these parameters (interactions are drawn independently except for the symmetry imposed by $\gamma$)
- when the simulation reaches equilibrium, we compute and store the values of $\langle\eta\rangle$, $\text{var}(\eta)$, and $\phi$ the fraction of survivors (species with $\eta_i > 0$)
- we create a smoothed and higher resolution map of $(\langle\eta\rangle, \text{var}(\eta), \phi)$ as a function of $(\bar{\beta}, \text{var}(\beta), \gamma)$, by using a Kernel Ridge Regression (package scikit-learn in Python, using KernelRidge for the regression and GridSearchCV to search for optimal regression parameters)
- for each dataset where we we know the observed relative yields at equilibrium $\eta_i$, we compute the Mahalanobis distance between the vectors $(\langle\eta\rangle, \text{var}(\eta), \phi)$ in the observed equilibrium and at each point of the map created above
- we select the parameter values which minimize this distance.

In general, this leads to a small (10-20%) change of $\bar{\beta}$ compared to the naive approximation, which improves our results somewhat. We also confirm the appropriateness of $\gamma \approx 0$ (e.g. for Wageningen we find an optimum at $\gamma \approx 0.16$, where the precise value of $\gamma$ has only a weak impact on the distance).

#### E.4 Metrics of agreement

Many different metrics may in principle be used to quantify the agreement between empirical coefficients and theoretical expectations, including regressions and comparisons to randomized data or other null models.

We found that each metric that we considered to estimate the agreement between predictions and observations could be misleading when taken on its own: it was possible to construct simulated interaction networks that would lead to high agreement scores, despite being visually starkly different from the theoretical pattern. Hence we rather construct a series of metrics, each focusing on a distinct feature, and our confidence in our conclusions relies only onwide agreement between the distributions of values obtained from empirical matrices, and those obtained from simulated matrices generated according to our theory.

Each metric $\rho_M$ is constructed by combining the following options:

- Which observed coefficients $z$ and theoretical expectations $E[z]$ are used:
  - for the pattern of means: empirical interactions $\beta_{ij}$ and their theoretical expectations $E[\beta_{ij}|\eta_i, \eta_j]$
  - for the pattern of correlations: the coefficients $y_{ijk}$ defined as

$$y_{ijk} = \frac{d_{ij} d_{ik}}{\sum_{m,n} d_{mn}^2} \qquad \text{where} \qquad d_{ij} = \beta_{ij} - E[\beta_{ij}|\eta_i, \eta_j] \qquad \text{(Eq. S35)}$$

    and their theoretical expectations i.e. the correlations $\mathrm{corr}(\beta_{ij}, \beta_{ik}|\eta_i, \eta_j, \eta_k)$.

- How the coefficients are binned:
  - By species $i$ (the affected species), or by the affecting species ($j$ or the $j, k$ pair)
  - In regular bins of $\eta_i$, or in bins of $\eta_j$ or $\eta_j \eta_k$
  - In regular bins of $E[z]$ (i.e. we group observed coefficients by what we predict to be their theoretical expectation, as in Fig. S1)

- The quantity $M$ that we compute within each bin of coefficients $z$:
  - the bin average
  - the $\eta$-dependence of coefficients within the bin (if coefficients are grouped by species $i$, their regression slope against $\eta_j$ or $\eta_j \eta_k$, and vice versa), see e.g. Fig S9

- How we compute the score $\rho_M$ of agreement between the empirical values $M_{emp}$ and the theoretical values $M_{theo}$ computed for each bin
  - The two-sided regression

$$\rho_M = \mathrm{cov}(M_{emp}, M_{theo}) / \max\left(\mathrm{var}(M_{emp}), \mathrm{var}(M_{theo})\right) \qquad \text{(Eq. S36)}$$

    which coincides with the Pearson correlation coefficient when $M_{emp}$ and $M_{theo}$ have the same variance, but decreases when their variances differ, so as to capture quantitative agreement and not only correlation[3].
  - The log-likelihood of $M$ given the theoretical mean $E[M]$ and variance $\mathrm{var}(M)$, i.e.

$$\rho_M = -\sum_{bins} (M_{emp} - E[N])^2 / 2\mathrm{var}(M_{theo}) \qquad \text{(Eq. S37)}$$

[3] We choose this definition over more standard alternatives because:

* a one-sided linear regression of $M_{emp}$ against $M_{theo}$ seems to overestimate agreement, finding slopes close to 1 even for experiments and simulations where the relationship is very nonlinear
* requiring only the correct ordering (by a Spearman rather than Pearson correlation) again tends to overestimate agreement between empirics and theory
* a correlation is normalized, whereas a distance between the two matrices would need some normalization (e.g. by comparison to some null model) to allow us to rank various experiments with different numbers of species along the same axis of agreement. This question of normalization introduces additional problems, such as what is the correct null model.

The first option puts more weight on the trends of $\beta$ with $\eta$, the last option puts more weight on the variance of $\beta$ around that trend: if the theoretical variance is large compared to the trend, observations with the same variance will achieve a good score in log-likelihood even if their trend is bad.)

In theory, these options and many others are all correct ways of assessing agreement between theory and data, due to the property of *self-averaging* exhibited by random networks (see Sec. B.2). This property means that there are infinitely many quantities that can be computed by some averaging over multiple coefficients in a given network, and, as we consider larger and larger network size $S$, *all* these quantities will eventually converge to their theoretical expectations in the ensemble of possible networks. In practice, some choices are more robust than others to the paucity of data (see Fig. S10).

#### E.5 Illustrations

##### E.5.1 Example 1: Coefficients binned by $\eta_i$

We show here how we compute agreement when coefficients are grouped in regular bins of $\eta_i$.

**Means**

- We are given one inferred matrix $\beta$ and a set of vectors $\boldsymbol{\eta}$ for the maximum diversity plots.
- If we have multiple abundance vectors $\boldsymbol{\eta}$, we may use bootstrapping to generate a much larger set (more precisely: we create more vectors by drawing $\eta_i$ at random from values for species $i$ in different plots).
- We take all triplets $(\eta_i, \eta_j, \beta_{ij})$, and bin them by $\eta_i$
- Within each bin, we perform a least squares fit of the constants $A$ and $B$ in

$$y = Ax + B, \qquad x = \eta_j, \qquad y = \beta_{ij} \qquad \text{(Eq. S38)}$$

- We finally compare $A$ to the theoretical prediction (Eq. S26) where $\eta_i$ is replaced by the bin average $\langle \eta_i \rangle_{bin}$

**Correlations**

- We take all quintuplets $(\eta_i, \eta_j, \eta_k, \beta_{ij}, \beta_{ik})$, and bin them by $\eta_i$
- Within each bin, we perform a least squares fit of the constant $C$ in

$$y = \sigma_\beta^2(\delta_{jk} + Cx), \qquad x = \eta_j \eta_k, \qquad y = (\beta_{ij} - \hat{\beta}_{ij})(\beta_{ik} - \hat{\beta}_{ik}) \qquad \text{(Eq. S39)}$$

  where $\delta_{jk} = 1$ if $j = k$, 0 otherwise, and $\sigma_\beta$ was inferred as noted above in Sec. E.3.1 (NB: it is important to keep diagonal terms $j = k$, since they contain part of the correlation; otherwise results worsen, as we checked on simulated data)

##### E.5.2 Example 2: Running averages

We show in Fig. S10 the process of constructing the empirical average matrix for the Wageningen experiment as shown in the main text, using the running average of coefficients $\beta_{ij}$ binned by their theoretical expectations. Fig. S11 shows the corresponding results for an assembled community of $S = 25$ species, starting from a random species pool.

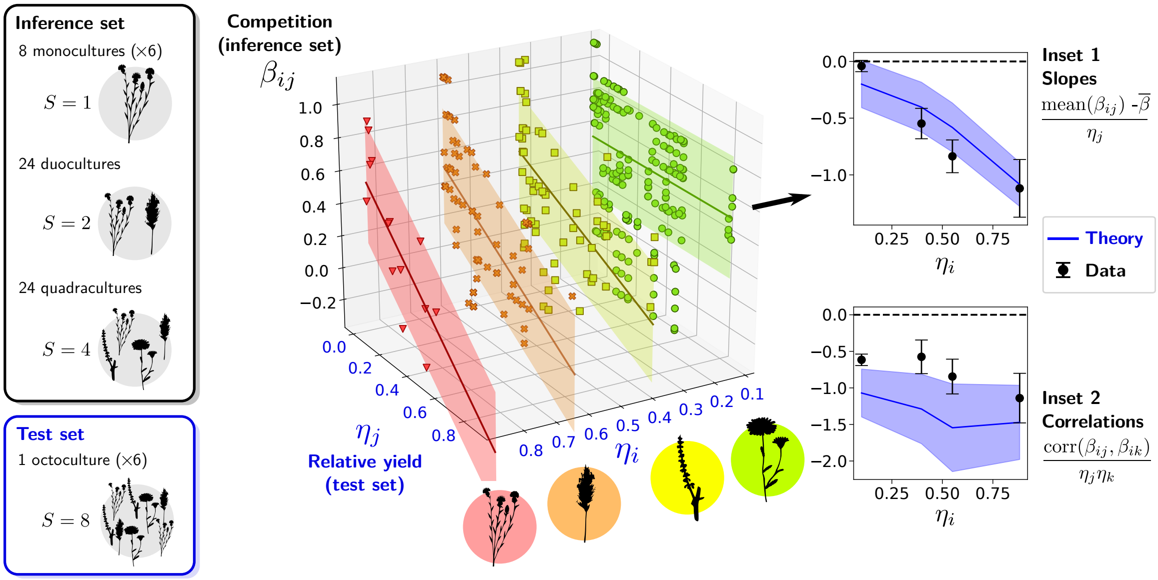

Figure S9: Visualization of the theoretical pattern in the Wageningen grassland experiment, when binning coefficients by $\eta_i$ (see Sec. E.5.1). We assume an equilibrium Lotka-Volterra description in which relative yields are given by $\eta_i(\boldsymbol{\theta}) = \theta_i$. In the theoretical pattern (Eq. S7), interactions $\beta_{ij}$ between two core species $i$ and $j$ (both with high relative yield $\eta$) are less competitive than average. Symbols represent triplets $(\eta_i, \eta_j, \beta_{ij})$ in the dataset (each point is a replica), grouped in four bins of $\eta_i$. (For legibility, we replace here the coordinate $\eta_i$ of each symbol by the bin's center of mass). For each bin, solid lines are linear fits of $\beta_{ij}$ against $\eta_j$, while the filled area indicates the standard deviation of $\beta_{ij}$ within each bin. Inset 1 shows the predicted and measured slopes of these lines. Inset 2 shows the anticorrelation pattern (Eq. S9), demonstrating that core species avoid competing against the same opponents. In both cases, theoretical predictions have no fitting parameter and are deduced solely from measured $\bar{\beta}$ and $\eta_i$. The filled region shows the theoretical range given the uncertainty on $\eta$, while dots represent measurements on data, with whiskers showing standard error on the mean.

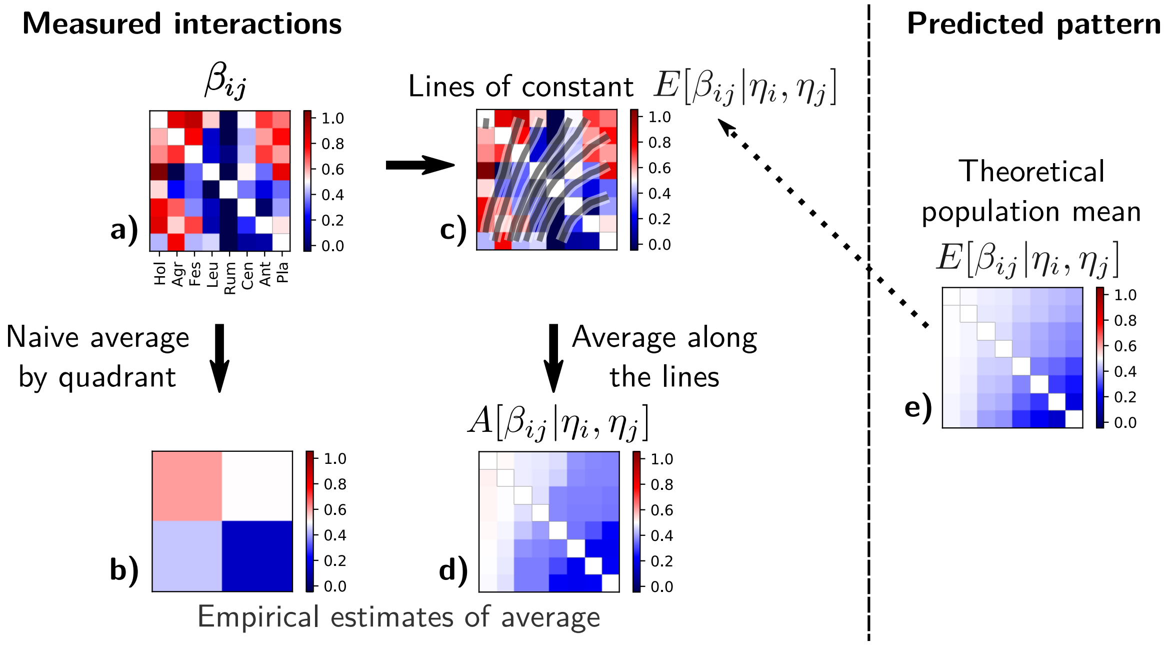

Figure S10: Comparison of measured interactions (**a**) and predicted pattern of means (**e**). We must perform some empirical average of coefficients $\beta_{ij}$, to compare to theoretical expectations. We show here a naive average by quadrants (**b**). We also show how to obtain the more robust running average employed in the main text: we make bins of $\beta_{ij}$, grouping together coefficients that have the same theoretical expectation $E[\beta_{ij}|\eta_i, \eta_j]$ (more precisely, we perform a running average of $\beta_{ij}$ versus $E[\beta_{ij}]$, see details in the main text). These bins correspond to level lines, i.e. lines of constant color, in the matrix (**e**). In (**c**), we superimpose these level lines over the measured matrix $\beta_{ij}$. Averaging together all the coefficients $\beta_{ij}$ that fall under the same line gives our empirical average **d**). In particular, we see that these level lines suggest to average together points that are in the top-left corner and middle-right of the same matrix, and hence the top-left corner should exhibit low average competition, rather than the high value found by chance in the naive quadrant method. This averaging procedure allows, with less data, to obtain better estimates than a naive smoothing of the matrix $\beta_{ij}$.

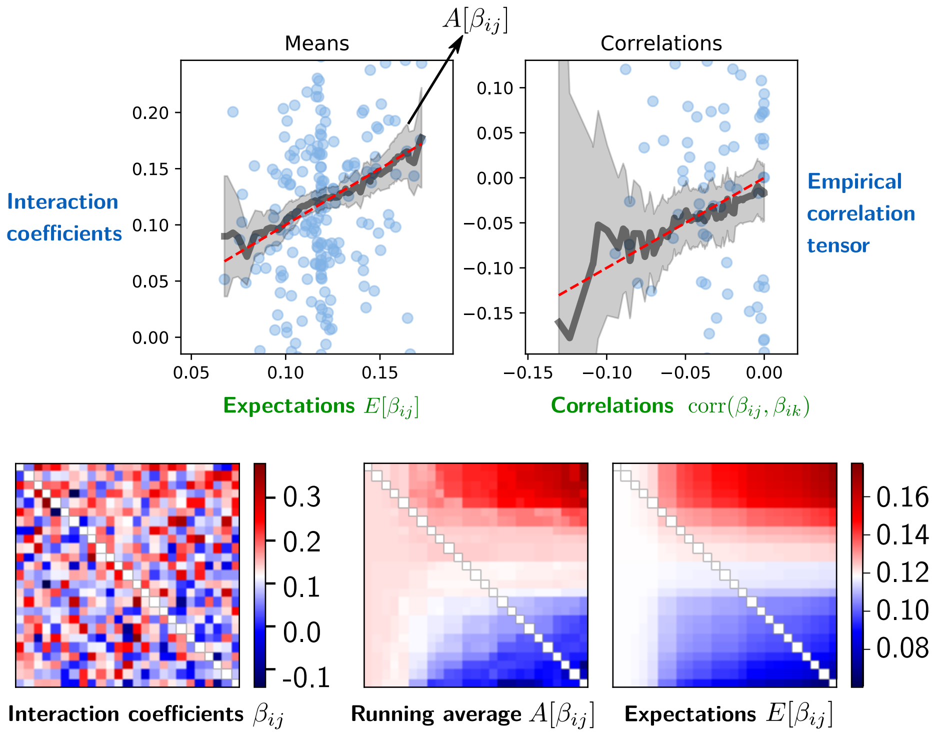

Figure S11: Results for simulated ecological assembly with $S = 25$ coexisting species (see protocol in [**?**]). The top row shows the predictions (Eq. S7) (left panel) and (Eq. S9) (right panel) on the x-axis, and the corresponding empirical estimates $A[\beta_{ij}]$ or $A[y_{ijk}]$ (as defined in Sec. A.2) on the y-axis. The symbols correspond to matrix elements (only a random subset is shown), and we plot the running average (grey curve, 90% CI in shaded area). The bottom row shows a matrix representation of interactions: the true interaction matrix $\beta_{ij}$, and both the empirical estimate $A[\beta_{ij}]$ and the theoretical prediction $E[\beta_{ij}]$ for the pattern of means.

### F Results

#### F.1 Scores for all experiments and simulations

In the main text, we do not directly show the metrics listed in Sec. E.4. These metrics cannot be compared directly between experiments, because each experiment has different equilibrium relative yields $\eta_i$ and a different mean and variance of interactions, $\bar{\beta}$ and $\mathrm{var}(\beta)$. This entails different theoretically-predicted ranges for each metric, as we show for multiple metrics and experiments in Fig. S12.

The metrics shown in Figures. S12, S13 and S14 are named following the explanations in Sec. E.4):

- `•/corr`: if corr, measurement for the pattern of correlations rather than the pattern of means
- `column/row`: binning coefficients by column or row
- `mean/slope`: taking the mean or linear regression slope within each bin
- `•/LL`: if LL, log-likelihood rather than two-sided regression

and the additional special metrics:

- `reldist`: two-sided regression between the running average of coefficients and their theoretical expectation
- `bilinear`: the overall bilinear slope of $\beta_{ij}$ with $\eta_i \eta_j$
- `smoothness`: we measure differences between adjacent coefficients, and compute the log of the fraction of these differences that are smaller than half the standard deviation of the whole matrix (a score of 0 would indicate perfect smoothness)
- `overall LL`: overall log-likelihood of elements $\beta_{ij}$ assuming that they were drawn from a multivariate normal distribution with the theoretical expectations (Eq. S7) and correlations (Eq. S9)

For each of these metrics, we computed the distribution of values obtained from bootstrapped empirical matrices, and found its mean and variance $\mu_{exp}$, $\sigma^2_{exp}$. We also computed the distribution of values in matrices generated according to our theory ($\mu_{th}$, $\sigma^2_{th}$), and in matrices generated with a competition-colonization trade-off ($\mu_{cc}$, $\sigma^2_{cc}$), with the same equilibrium as the data. We define similarity as

$$s = \begin{cases} 1 & \text{if} \quad |\mu_{exp} - \mu_{th}| < 2\sigma_{th} \\ -1 & \text{otherwise} \end{cases} \qquad \text{(Eq. S40)}$$

which states in a simple binary score whether observed values fall within the confidence interval (2 standard deviations) of predicted values. We define discrimination as

$$D = \left( 1 + \min\left( \frac{|\mu_{exp} - \mu_{th}|}{\sigma_{exp} + \sigma_{th}}, \frac{|\mu_{cc} - \mu_{th}|}{\sigma_{cc} + \sigma_{th}} \right)^{-1} \right)^{-1} \qquad \text{(Eq. S41)}$$

which quantifies how well a given metric can differentiate between patterns ($D$ goes to 0 when standard deviations are significantly larger than the interval between means). The product $s \times D$ gives our final score.

We show these scores for our set of metrics and for all experiments in Fig. S13 and S14. As additional illustrations, we also show results for four 8-species simulated matrices following our pattern (`simugaussS8`) and four 16-species simulated matrices following a competition-colonization trade-off pattern (`simuccoptS16`). In the case of the matrices following our pattern, we further impose that they have the same equilibrium relative yields $\eta_i$ as the Wageningen experiment, and the same average and variance of interaction strengths.

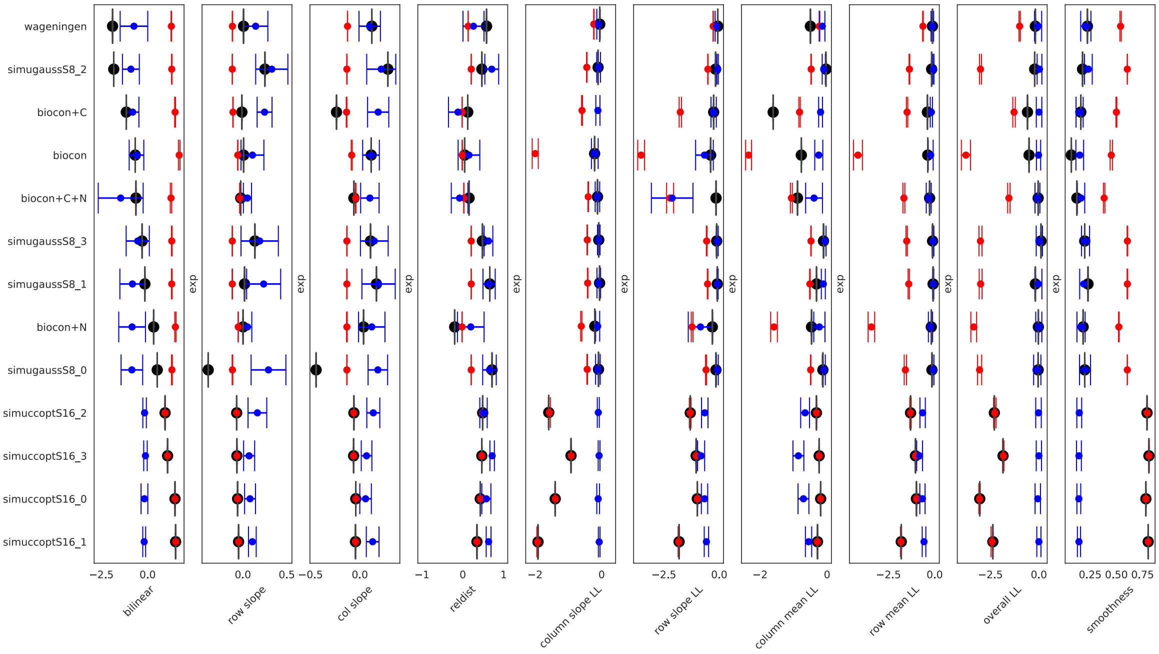

Figure S12: For multiple metrics pertaining to the pattern of means (defined in Sec. F.1), we show: in blue, the predicted range of values for this metric in our theoretical pattern (90% CI of values obtained in simulated matrices generated with our theory); in red, the predicted range of values for some counter-examples to our theory (90% CI for matrices generated with a colonization-competition trade-off); as a large black dot, the observed value of the metric in the median empirical matrix (i.e. the matrix whose elements are the median of all the values obtained for each $\beta_{ij}$ in our different bootstrapped replicas, see Appendix A). We can see that the expected range of values for each metric depends on the experiment. Thus, we conclude on agreement between theory and data only if: the black dot falls in the blue range (similarity score defined in Sec. F.1), and the blue and red range are well-separated (discrimination score).

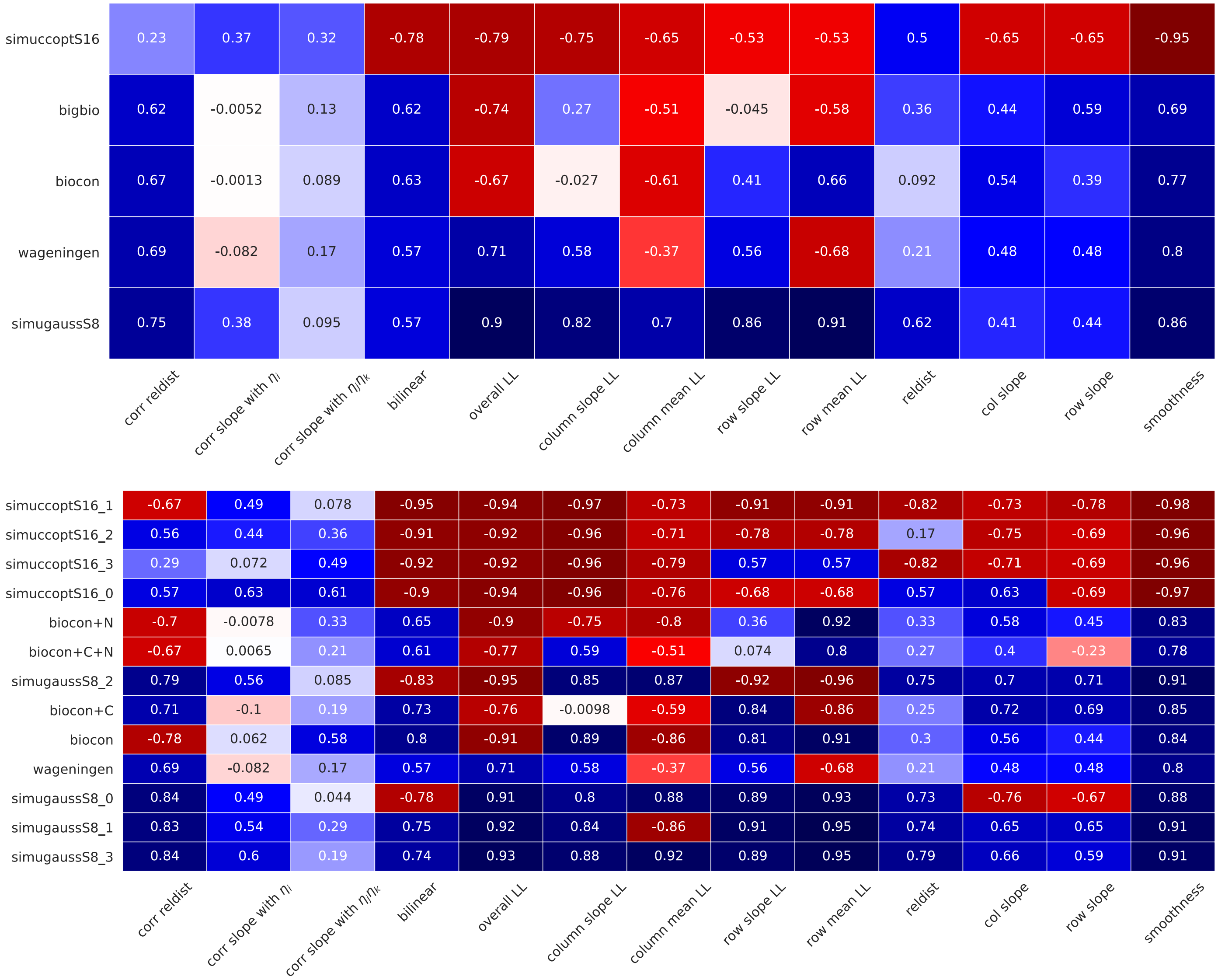

Figure S13: Comparison of metrics of agreement between predicted and measured mean interaction pattern (defined in Sec. E.4 and Appendix F) for all bootstrap replicas of all experiments and simulations studied in this work, grouping together various experimental treatments or simulated replicates (top). Details per treatment for all experiments except the Big Bio experiment shown in Fig. S14 (bottom).

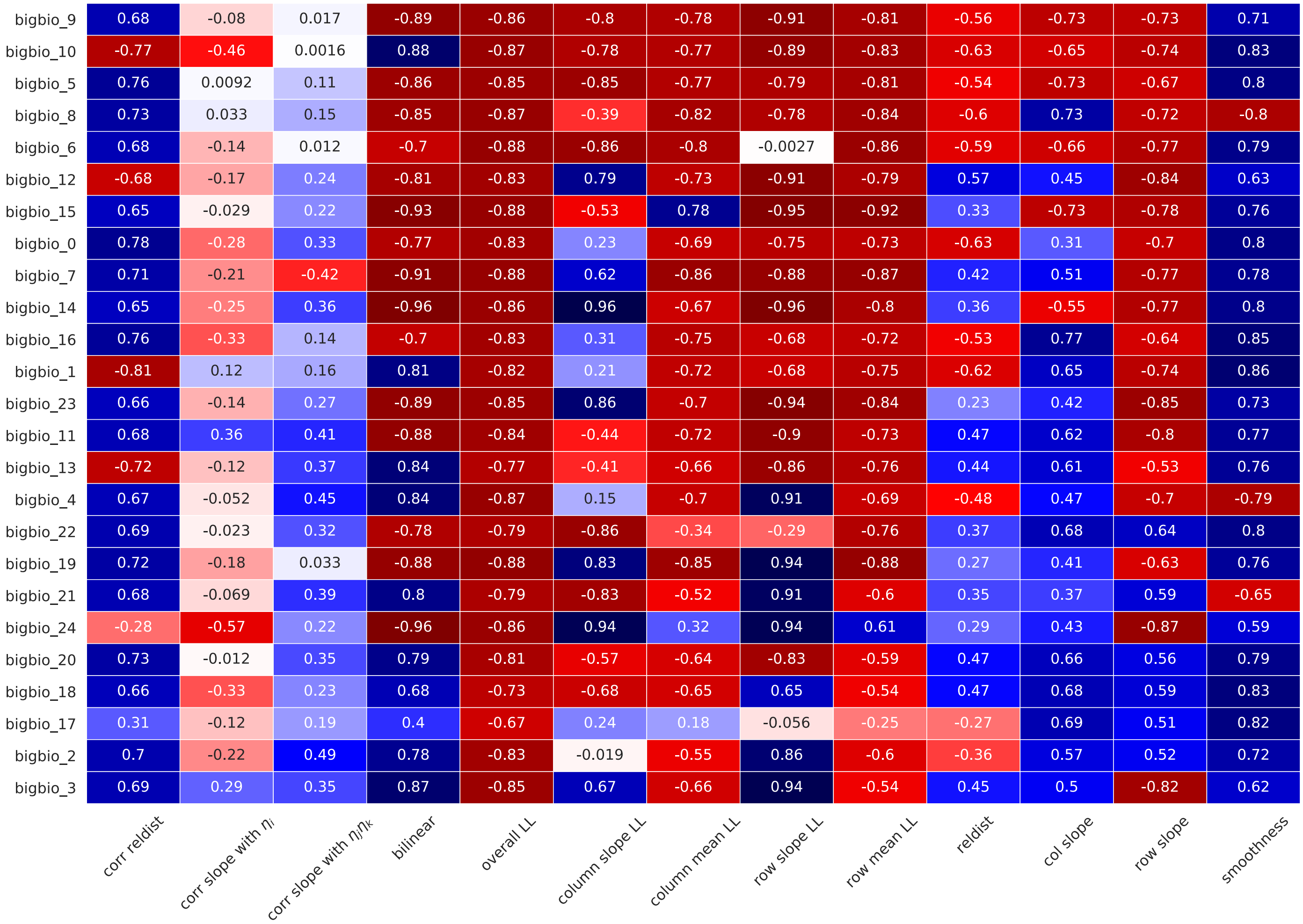

Figure S14: Comparison of metrics of agreement between predicted and measured mean interaction pattern (defined in Sec. E.4 and Appendix F) for all bootstrap replicas of all 24 compositions with more than 8 species that we retained and tested in the Big Bio experiment.

#### F.2 Per-species results for all experiments

This final section displays more in-detail graphs for the experimental data (Figs. S15 to S17), focusing on the pattern of means and per-species binning, to help visualize and estimate more intuitively the agreement between predictions and data, and what this agreement means for particular species.

We show here the inferred matrix $\beta_{ij}$ where missing elements are those whose variance between bootstrapped replicas (see Appendix A) exceeded the variance between median elements in the matrix (indicating poor inference). We also show the theoretical prediction for expectation $E[\beta_{ij}|\eta_i, \eta_j]$, and the row-wise and column-wise means and slopes, following the conventions in Fig. 3 (main text): dots are empirical values, the solid line is the average over simulated matrices generated with our theory, the filled area includes $\pm 1$ standard deviation among simulated matrices, and the dashed lines represent the 90% confidence interval for these theoretical predictions.

##### F.2.1 Comments on particular experiments

We only show two representative examples out of the 24 tested compositions in the Big Bio experiment (entitled Cedarcreekcomp6 and Cedarcreekcomp16 below) as an illustration. We remark that inferred interaction coefficients $\beta_{ij}$ in the BioCON experiments have many missing elements (high variance between bootstrapped replicas), whereas fewer coefficients $\beta_{ij}$ are missing in the BigBio experiment, and the ones that we retain have a narrower distribution (still notably wider than the one found in the Wageningen experiment). We ascribe this difference to the absence of duocultures in the BioCON experiment, increasing the uncertainty on pairwise interactions, especially for species that are rare in polycultures with $S \geq 4$.

##### F.2.2 Comments on particular metrics

It appears that scores based on our correlation pattern $\text{corr}(\beta_{ij}, \beta_{ik})$ can be good even for matrices that have not been generated according to our theory (the `simuccoptS16` examples). Nevertheless, in most experiments, we can still use correlations to discriminate between an instance of our theory and a counter-example, as shown by sufficiently high discrimination scores.

Finally, we note that the "column mean" is the metric that is most systematically wrong and variable between species. We believe that this is an artefact of our inference method: when the carrying capacity of a species is wrongly inferred, we can systematically overestimate or underestimate the success of this species, and therefore make the opposite error on the interaction coefficients, which are defined as the impact of this species on others divided by this species' success.

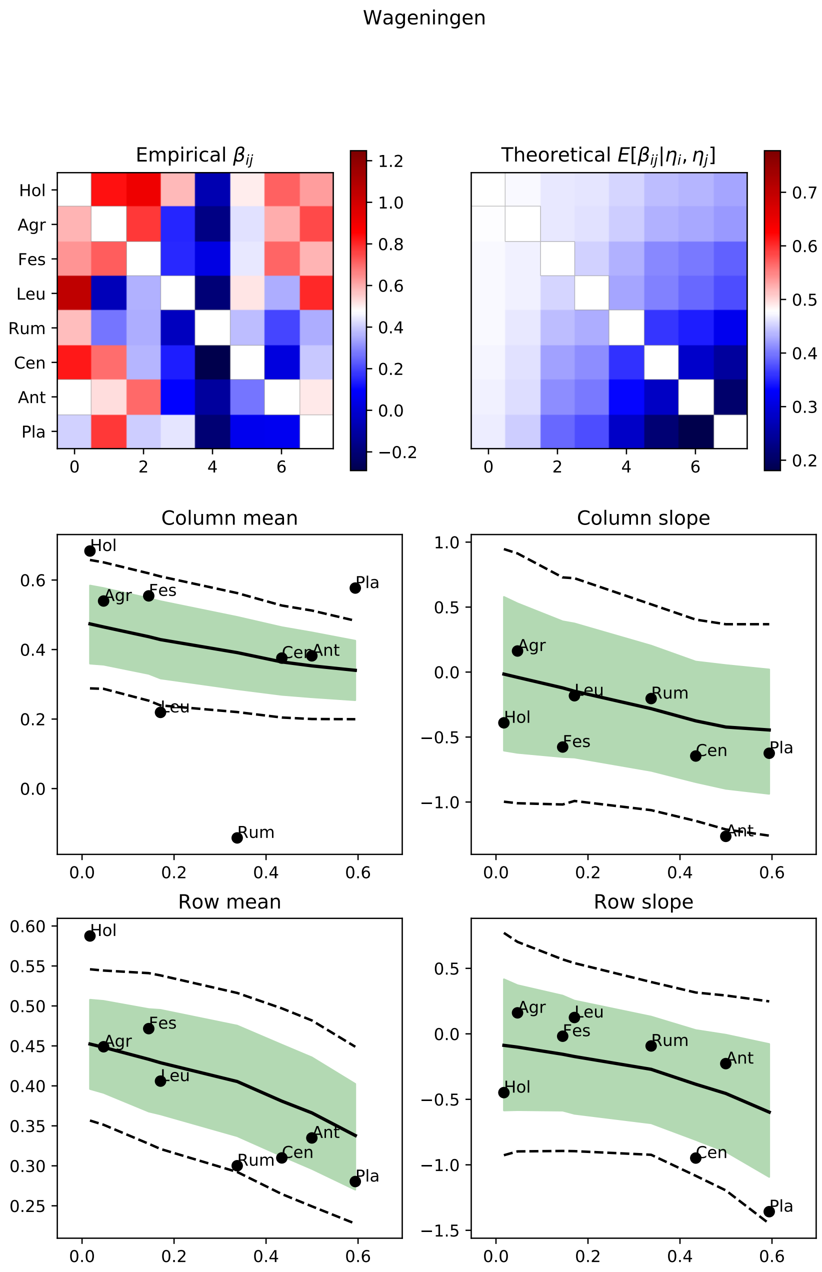

Figure S15: Detailed results for the Wageningen experiment.

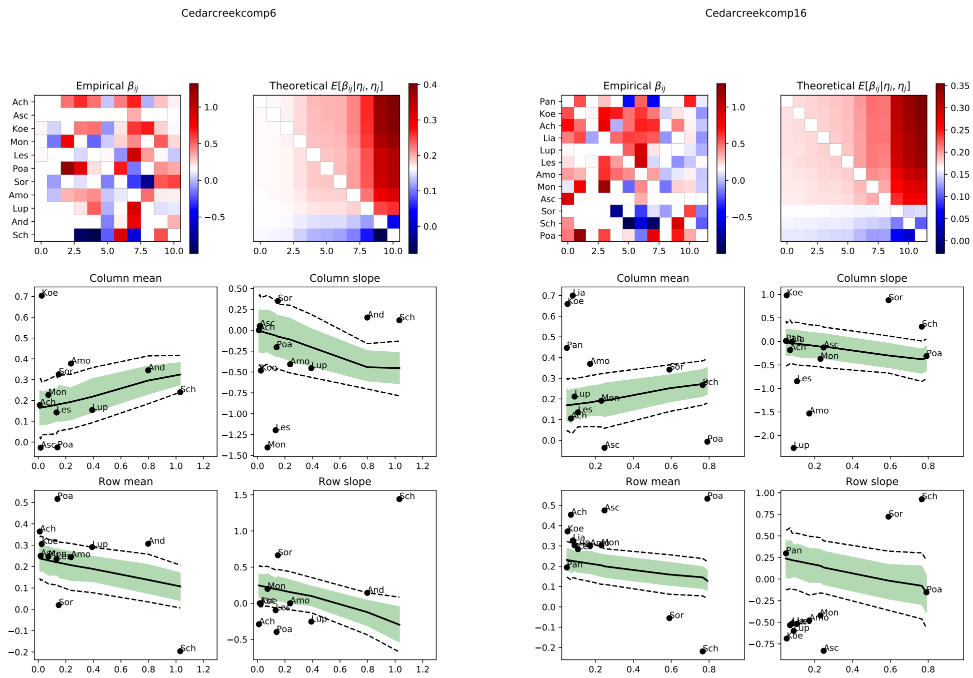

Figure S16: Detailed results for the Big Bio experiment (two compositions).

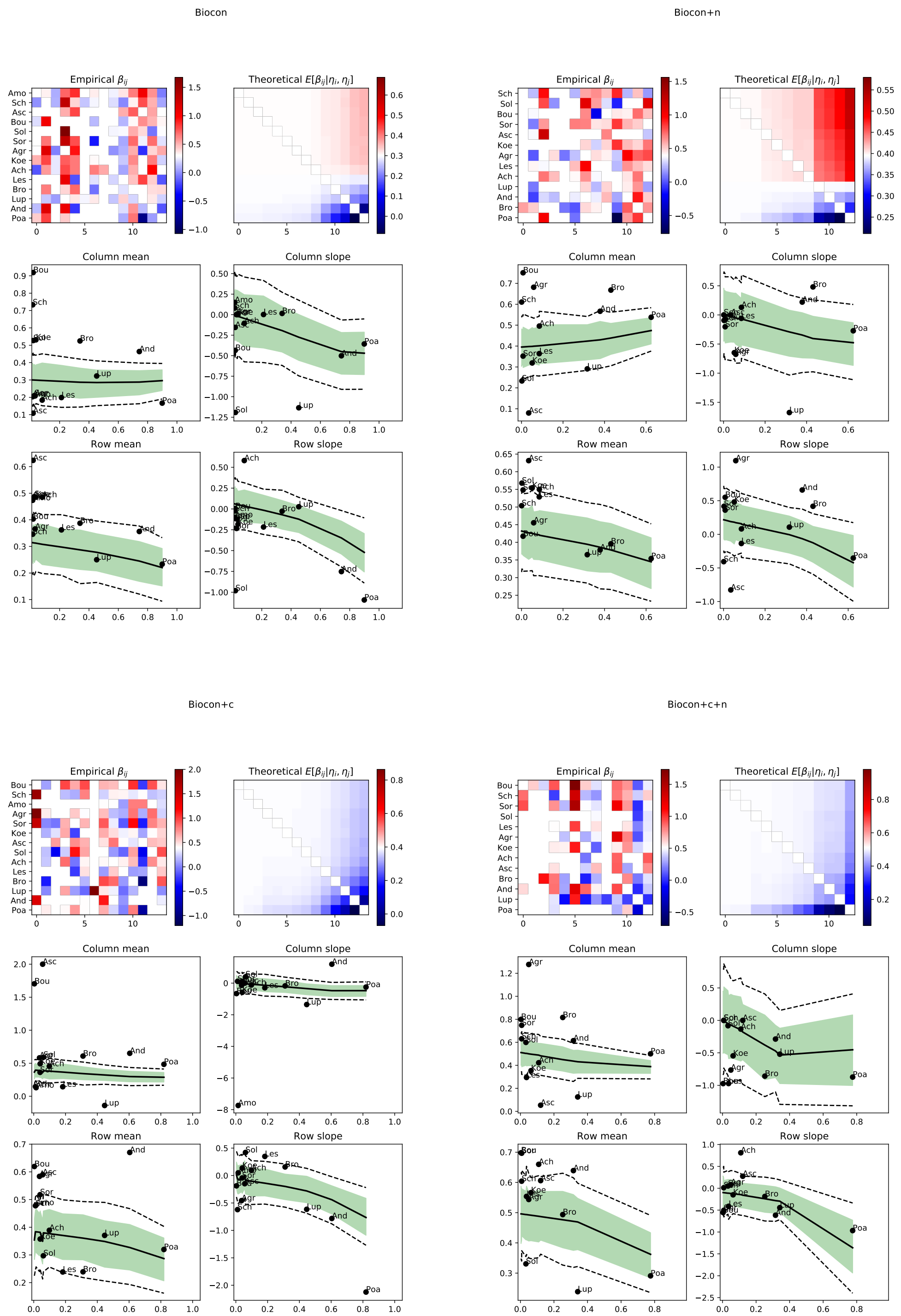

Figure S17: Detailed results for the BioCON results (four treatments: control, +N, +CO2, +N +CO2).
